# Meta-GWAS Accuracy and Power (MetaGAP) calculator shows that hiding heritability is partially due to imperfect genetic correlations across studies

**DOI:** 10.1101/048322

**Authors:** Ronald de Vlaming, Aysu Okbay, Cornelius A. Rietveld, Magnus Johannesson, Patrik K.E. Magnusson, André G. Uitterlinden, Frank J.A. van Rooij, Albert Hofman, Patrick J.F. Groenen, A. Roy Thurik, Philipp D. Koellinger

## Abstract

Large-scale genome-wide association results are typically obtained from a fixed-effects meta-analysis of GWAS summary statistics from multiple studies spanning different regions and/or time periods. This approach averages the estimated effects of genetic variants across studies. In case genetic effects are heterogeneous across studies, the statistical power of a GWAS and the predictive accuracy of polygenic scores are attenuated, contributing to the so-called ‘missing heritability’. Here, we describe the online Meta-GWAS Accuracy and Power calculator (MetaGAP; available at www.devlaming.eu) which quantifies this attenuation based on a novel multi-study framework. By means of simulation studies, we show that under a wide range of genetic architectures, the statistical power and predictive accuracy provided by this calculator are accurate. We compare the predictions from MetaGAP with actual results obtained in the GWAS literature. Specifically, we use genomic-relatedness-matrix restricted maximum likelihood (GREML) to estimate the SNP heritability and cross-study genetic correlation of height, BMI, years of education, and self-rated health in three large samples. These estimates are used as input parameters for the MetaGAP calculator. Results from the calculator suggest that cross-study heterogeneity has led to attenuation of statistical power and predictive accuracy in recent large-scale GWAS efforts on these traits (e.g., for years of education, we estimate a relative loss of 51–62% in the number of genome-wide significant loci and a relative loss in polygenic score *R*^2^ of 36–38%). Hence, cross-study heterogeneity contributes to the missing heritability.

**Author Summary:** Large-scale genome-wide association studies are uncovering the genetic architecture of traits which are affected by many genetic variants. Such studies typically meta-analyze association results from multiple studies spanning different regions and/or time periods. GWAS results do not yet capture a large share of the total proportion of trait variation attributable to genetic variation. The origins of this so-called ‘missing heritability’ have been strongly debated. One factor exacerbating the missing heritability is heterogeneity in the effects of genetic variants across studies. Its influence on statistical power to detect associated genetic variants and the accuracy of polygenic predictions is poorly understood. In the current study, we derive the precise effects of heterogeneity in genetic effects across studies on both the statistical power to detect associated genetic variants as well as the accuracy of polygenic predictions. We provide an online calculator, available at www.devlaming.eu, which accounts for these effects. By means of this calculator, we show that imperfect genetic correlations between studies substantially decrease statistical power and predictive accuracy and, thereby, contribute to the missing heritability. The MetaGAP calculator helps researchers to gauge how sensitive their results will be to heterogeneity in genetic effects across studies. If strong heterogeneity is expected, random-instead of fixed-effects meta-analysis methods should be used.

## Introduction

Large-scale GWAS efforts are rapidly elucidating the genetic architecture of polygenic traits, including anthropometrics [1, 2] and diseases [3–5], as well as behavioral and psychological outcomes [6–8]. These efforts have led to new biological insights, therapeutic targets, and polygenic scores (PGS), and help to understand the complex interplay between genes and environments in shaping individual outcomes [7, 9, 10]. However, GWAS results do not yet account for a large part of the estimated heritability [1, 2, 7, 8]. This dissonance, which is referred to as the ‘missing heritability’, has received broad attention [11–17]. Missing heritability can be divided into two parts: ‘still-missing heritability’ [15–17] – defined as the difference between the estimate of heritability based on family data (*h*^2^) and the SNP-based estimate 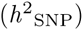, where 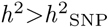 – and ‘hiding heritability’ [15–17] – defined as the difference between 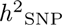 and the estimate based on genetic variants that reach genome-wide significance in a GWAS 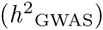, where 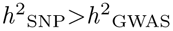.

Amongst others, four factors have been proposed to explain the missing heritability. First, standard genotyping techniques overlook some genetic variation explained by poorly tagged rare variants [18]. Second, non-additive genetic effects (e.g., epistasis) may inflate *h*^2^, creating so-called ‘phantom heritability’ [14]. Third, GWAS sample sizes are not large enough to fully capture 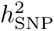 [18, 19]. Fourth, differences across strata (e.g., studies) in genetic effects, phenotype measurement, and phenotype accuracy lead to loss of signal [20–22], attenuating both the power of a GWAS [17, 20, 23, 24] and the predictive accuracy of the PGS in a hold-out sample [25]. We focus on the contribution of one such form of heterogeneity to missing heritability, viz., heterogeneity across studies.

Bearing the aforementioned attenuation of statistical power and PGS accuracy in mind, cross-study heterogeneity decreases the chances of a study to yield meaningful results [24, 26]. Therefore, the precise attenuation arising from such heterogeneity should be well understood. Nevertheless, a theoretical multi-study framework, relating statistical power and predictive accuracy to cross-study heterogeneity, is still absent. We bridge this gap by developing a Meta-GWAS Accuracy and Power (MetaGAP) calculator (available at www.devlaming.eu) that accounts for the cross-study genetic correlation (CGR). This calculator infers the statistical power to detect associated SNPs and the predictive accuracy of the PGS in a meta-analysis of GWAS results from genetically and phenotypically heterogeneous studies. Using simulations, we show that the MetaGAP calculator is accurate under a wide range of genetic architectures, even when the assumptions of the calculator are violated.

Although meta-analysis methods accounting for heterogeneity exist [27–31], large-scale GWAS results are typically still obtained from fixed-effects meta-analysis methods [32, 33] such as implemented in `METAL` [34]. Therefore, it is important to infer the attenuation in statistical power and PGS accuracy when applying such a fixed-effects meta-analysis method. Hence, the MetaGAP calculator assumes the use of a fixed-effects meta-analysis method. Consequently, the calculator will help researchers to assess the merits of an intended fixed-effects meta-analysis of GWAS results and to gauge whether it is more appropriate to apply a meta-analysis method that accounts for heterogeneity.

Since heterogeneity can lead to differences in heritability between studies, this raises the important question which ‘target heritability’ we are referring to, when considering the proportion of hiding heritability explained by heterogeneity. For statistical power, we define the target heritability as the sample-size weighted average of the within-study estimates of SNP heritability for the studies included in the meta-analysis. For PGS accuracy, we set SNP heritability in the hold-out study as target. Under these definitions, we posit that the expected statistical power and PGS accuracy in the presence of heterogeneity fall short of what is expected under homogeneity.

In an empirical application, we use genomic-relatedness-matrix restricted maximum likelihood (GREML) to estimate the SNP-based heritability and CGR of several polygenic traits across three distinct studies: the Rotterdam Study (RS), the Swedish Twin Registry (STR), and the Health and Retirement Study (HRS). For self-rated health, years of education, BMI, and height, we obtain point-estimates of CGR between 0.47 and 0.97, suggesting that even extremely large GWAS meta-analyses will fall short of explaining the full 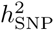 for these traits. We use the estimates of SNP heritability and CGR to quantify the expected number of hits and the predictive accuracy of the PGS in recent GWAS efforts for these traits. Our theoretical predictions align with empirical observations. By comparing these figures to the predicted number of hits and PGS accuracy under perfect CGRs, we show that imperfect CGRs lead to considerable attenuation of both (e.g., for height under an estimated CGR of 0.97, the expected relative loss in the number of hits is 8–9% and the relative loss in PGS *R*^2^ is 6–7%, whereas for years of education under an estimated CGR of 0.78, we expect a relative loss of 51–62% in the number of genome-wide significant loci and a relative loss in polygenic score *R*^2^ of 36–38%). Hence, heterogeneity can explain a considerable part of the hiding heritability.

## Materials and Methods

### Definitions and assumptions

The MetaGAP calculator is based on theoretical expressions for statistical power and PGS accuracy, derived in S1 Derivations Power and S2 Derivations Accuracy. In these expressions, within-study estimates of SNP heritability (e.g., inferred using `GCTA` [35]) are important input parameters. Estimates of CGR (e.g., inferred as genetic correlations across studies using pairwise bivariate methods implemented in `GCTA` [35] and LD-score regression [36]) also play a central role in those expressions. Importantly, as we show in S3 Note on Genetic Correlations, such estimates of CGR are affected by the cross-study overlap in trait-affecting loci as well as the cross-study correlation in the effects of these overlapping loci. In our derivations, we assume that the set of trait-affecting loci is the same across all studies and that, consequently, CGRs are shaped solely by cross-study correlations in the effects. Using simulation studies, discussed in S4 Simulation Studies, we assess how violations of this assumption affect our results.

In line with other work, we define the effective number of SNPs, *S*, as the number of haplotype blocks (i.e., independent chromosome segments) [37], where variation in each block is tagged by precisely one genotyped SNP. By genotyped SNPs we also mean imputed SNPs. Hence, in our framework, there are *S* SNPs contributing to the polygenic score. Due to linkage disequilibrium (LD) this number is likely to be substantially lower than the total number of SNPs in the genome [38], and is inferred to lie between as little as 60,000 [15] and as much as 5 million [38].

In terms of trait-affecting variants, we consider a subset of *M* SNPs from the set of *S* SNPs. Each SNP in this subset tags variation in a segment that bears a causal influence on the phenotype. We refer to *M* as the associated number of SNPs. We assume that the *M* associated SNPs jointly capture the full SNP-based heritability for the trait of interest and, moreover, that each associated SNP has the same theoretical *R*^2^ with respect to the phenotype. In the simulation studies, we also assess the impact of violations of this ‘equal-*R*^2^’ assumption.

By considering only independent genotyped SNPs that are assumed to fully tag the causal variants, we can ignore LD among genotyped variants and between the causal variant and the genotyped variants. Thereby, we can greatly reduce the theoretical and numerical complexity of the MetaGAP calculator. However, a genotyped tag SNP does not necessarily capture the full variation of the causal variant present in that independent segment. Nevertheless, the inputs for SNP heritability used in the MetaGAP calculator are within-study GREML estimates of heritability, based on the available (common) SNPs. Therefore, if these genotyped SNPs are in imperfect LD with the causal variants, this will lead to a downward bias in the SNP-based heritability estimates [39]. Hence, the imperfect tagging of the causal variants is already absorbed by downward bias in the SNP-based heritability estimates.

### Power of a GWAS meta-analysis under heterogeneity

The theoretical distribution of the *Z* statistic, resulting from a meta-analysis of GWAS results under imperfect CGRs, can be found in S1 Derivations Power. These expressions allow for differences in sample size, 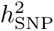, and CGR across (pairs of) studies. For intuition, we here present the specific case of a meta-analysis of results from two studies with CGR *ρ*_G_, with equal SNP-based heritability 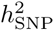, and equal sample sizes (i.e., *N* in Study 1 and *N* in Study 2). Under this scenario, we find that under high polygenicity, the *Z* statistic of an associated SNP *k* is normally distributed with mean zero and the following variance:

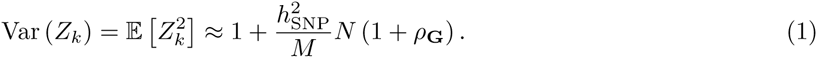

Fixed-effects meta-analysis approaches are still frequently used in large-scale GWAS efforts. Therefore, we consider statistical power when applying this type of meta-analysis, while assuming that the actual data-generating process follows a random-effects model, where cross-study correlations in SNP effects shape the inferred CGRs. When one has random effects, under the null hypothesis a SNP effect follows a degenerate distribution with all probability mass at zero, whereas under the alternative hypothesis a SNP effect follows a distribution with mean zero and a finite non-zero variance. Bearing in mind that we can write a meta-analysis *Z* statistic as weighted a average of true effects across studies and noise terms, the null hypothesis leads to a *Z* statistic with a mean equal to zero and a variance equal to one, whereas the alternative hypothesis does not lead to a non-zero mean in the *Z* statistic, but rather to excess variation (i.e., a variance larger than one).

The larger the variance in the *Z* statistic, the higher the probability of rejecting the null. The ratio of 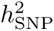 and *M* can be regarded as the theoretical *R*^2^ of each associated SNP with respect to the phenotype. Eq. 1 reveals that (i) when sample size increases, power increases, (ii) when 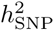 increases, the *R*^2^ per associated SNP increases and therefore power increases, (iii) when the number of associated SNPs increases, the *R*^2^ per associated SNP decreases and therefore power decreases, (iv) when the CGR is zero the power of the meta-analysis is identical to the power obtained in each of the two studies when analyzed separately, yielding no strict advantage to meta-analyzing, and (v) when the CGR is plus one the additional variance in the *Z* statistic relatively to the variance under the null is twice the additional variance one would have when analyzing the studies separately, yielding a strong advantage to meta-analyzing.

Notably, our expression for 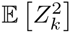 bears a great resemblance to expressions for the expected value of the squared *Z* statistic when accounting for LD, population stratification, and polygenicity [36, 40, 41]. Consider the scenario where the CGR equals one between two samples of equal size. Based of Eq. 1, we then have that 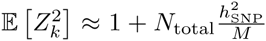 for a trait-affecting haplotype block, where *N*_total_ = 2*N*. This expression is equivalent to the expected squared *Z* statistic from the linear regression analysis for a trait-affecting variant reported in Section 4.2 of the Supplementary Note to [41] as well as the first equation in [36] when assuming that confounding biases and LD are absent.

In order to compute statistical power in a multi-study setting, we first use the generic expression for the variance of the GWAS *Z* statistic derived in S1 Derivations Power to characterize the distribution of the *Z* statistic under the alternative hypothesis. Given a genome-wide significance threshold (denoted by *α*; usually *α* = 5 · 10^−8^), we use the normal cumulative distribution function under the alternative hypothesis to quantify the probability of attaining genome-wide significance for an associated SNP. This probability we refer to as the ‘power per associated SNP’ (denoted here by *β*). Given that we use SNPs tagging independent haplotype blocks, we can calculate the probability of rejecting the null for at least one SNP and the expected number of hits, true positives, false positives, false negatives, and positive negatives, as functions of *α*, *β*, the number of truly associated SNPs (denoted by *M*), and the number of non-associated SNPs (denoted by *S* − *M*). Specifically,

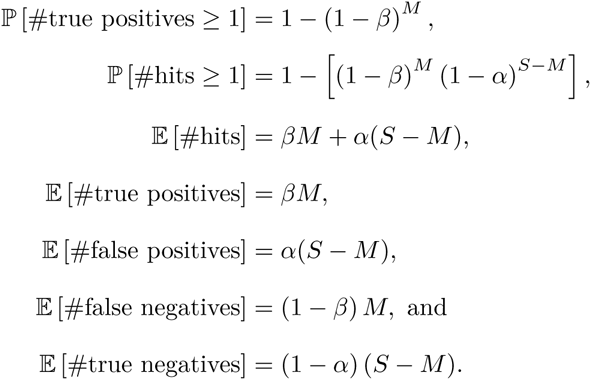

### *R*^2^ of a polygenic score under heterogeneity

In S2 Derivations Accuracy we derive a generic expression for the theoretical *R*^2^ of a PGS in a hold-out sample, with SNP weights based on a meta-analysis of GWAS results under imperfect CGRs. We consider a PGS that includes all the SNPs that tag independent haplotype blocks (i.e., there is no SNP selection).

For intuition, we here present an approximation for prediction in a hold-out sample, with SNP weights based on a GWAS in a single discovery study with sample size *N*, where both studies have SNP heritability 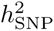, and with CGR *ρ*_*G*_, between the studies. Under high polygenicity, the *R*^2^ of the PGS in the hold-out sample is then given by the following expression:

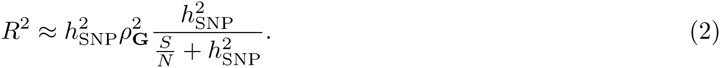

In case the CGR is one, and we consider the *R*^2^ between the PGS and the genetic value (i.e., the genetic component of the phenotype) instead of the phenotype itself, the first two terms in Eq. 2 disappear, yielding an expression equivalent to the first equation in [37]. Assuming a CGR of one and that all SNPs are associated, Eq. 2 is equivalent to the expression in [25] for the *R*^2^ between the PGS and the phenotype in the hold-out sample.

From Eq. 2, we deduce that (i) as the effective number of SNPs *S* increases, the *R*^2^ of the PGS deteriorates (since every SNP-effect estimate contains noise, owing to imperfect inferences in finite samples), (ii) given the effective number of SNPs, under a polygenic architecture, the precise fraction of effective SNPs that is associated does not affect the *R*^2^, (iii) *R*^2^ is quadratically proportional to *ρ*_*G*_, implying a strong sensitivity to CGR, and (iv) as the sample size of the discovery study grows, the upper limit of the *R*^2^ is given by 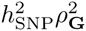, implying that the full SNP heritability in the hold-out sample cannot be entirely captured as long as CGR is imperfect.

### Online power and *R*^2^ calculator

An online version of the MetaGAP calculator can be found at www.devlaming.eu. This calculator computes the theoretical power per trait-affecting haplotype block, the power to detect at least one of these blocks, and the expected number of (a) independent hits, (b) true positives, (c) false positives, (d) false negatives, and (e) true negatives, for a meta-analysis of GWAS results from *C* studies. In addition, it provides the expected *R*^2^ of a PGS for a hold-out sample, including all GWAS SNPs, with SNP weights based on the meta-analysis of the GWAS results from *C* studies. Calculations are based on the generic expressions for GWAS power derived in S1 Derivations Power and PGS *R*^2^ derived in S2 Derivations Accuracy.

The calculator assumes a quantitative trait. Users need to specify either the average sample size per study or the sample size of each study separately. In addition, users need to specify either the average within-study SNP heritability or the SNP heritability per study. The SNP heritability in the hold-out sample also needs to be provided. Users are required to enter the effective number of causal SNPs and the effective number of SNPs in total. The calculator assumes a fixed CGR between all pairs of studies included in the meta-analysis and a fixed CGR between the hold-out sample and each study in the meta-analysis. Hence, one needs to specify two CGR values: one for the CGR within the set of meta-analysis studies and one to specify the genetic overlap between the hold-out sample and the meta-analysis studies.

Finally, a more general version of the MetaGAP calculator is provided in the form of `MATLAB` code (www.mathworks.com), also available at www.devlaming.eu. This code can be used in case one desires to specify a more versatile genetic-correlation matrix, where the CGR can differ between all pairs of studies. Therefore, this implementation requires the user to specify a full (*C*+1)-by-(*C*+1) correlation matrix. Calculations in this code are fully in line with the generic expressions in S1 Derivations Power and S2 Derivations Accuracy.

### Assessing validity of theoretical power and *R*^2^

We simulate data for a wide range of genetic architectures in order to assess the validity of our theoretical framework. As we show in S4 Simulation Studies, the theoretical expressions we derive for power and *R*^2^ are accurate, even for data generating processes substantially different from the process we assume in our derivations. Our strongest assumptions are that all truly associated SNPs have equal *R*^2^ with respect to the phenotype regardless of allele frequency and that genome-wide CGRs are shaped solely by the cross-study correlations in the effects of causal SNPs. When we simulate data where the former assumption fails and where – in addition – allele frequencies are non-uniformly distributed and different across studies, the root-mean-square prediction error of statistical power lies below 3% and that of PGS *R*^2^ below 2%. Moreover, when we simulate data where the CGR is shaped by both non-overlapping causal loci across studies and the correlation of the effects of the overlapping loci, the RMSE is less than 2% for both statistical power and PGS *R*^2^.

### Estimating SNP heritability and CGR

Using 1000 Genomes-imputed (1kG) data from the RS, STR, and HRS, we estimate SNP-based heritability and CGR respectively by means of univariate and bivariate GREML [35, 42] as implemented in `GCTA` [35]. In our analyses we consider the subset of HapMap3 SNPs available in the 1kG data. In S5 Data and Quality Control we report details on the genotype and phenotype data, as well as our quality control (QC) procedure. After QC we have a dataset, consisting of ≈ 1 million SNPs and ≈ 20,000 individuals, from which we infer 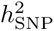 and CGR. In S6 GREML Estimation we provide details on the specifications of the models used for GREML estimation.

## Results

### Determinants of GWAS power and PGS *R*^2^

Using the MetaGAP calculator, we assessed the theoretical power of a meta-analysis of GWAS results from genetically heterogeneous studies and the theoretical *R*^2^ of the resulting PGS in a hold-out sample, for various numbers of studies and sample sizes, and different values of CGR and 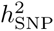.

#### Sample size and CGR

Fig. 1 shows contour plots for the power per truly associated SNP and *R*^2^, for a setting with 50 studies, for a trait with 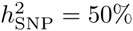, for various combinations of total sample size and CGR. Increasing total sample size enhances both power and *R*^2^. When the CGR is perfect, power and *R*^2^ (relative to 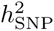) have a near-identical response to sample size. This similarity in response gets distorted when the CGR decreases. For instance, in the scenario of 100k SNPs of which a subset of 1k SNPs is causal with 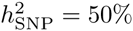, in a sample of 50 studies with a total sample size of 10 million individuals, a CGR of one yields 94% power per causal SNP and an *R*^2^ of 49%, which is 98% of the SNP heritability, whereas for a CGR of 0.2 the power is still 87% per SNP, while the *R*^2^ of the PGS is 8.5%, which is only 17% of 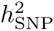. Thus, *R*^2^ is far more sensitive to an imperfect CGR than the meta-analytic power is. This finding is also supported by the approximations of power in in Eq. 1 and of PGS *R*^2^ in Eq. 2; these expressions show that, for two discovery studies, the CGR has a linear effect on the variance of the meta-analysis *Z* statistic, whereas, for one discovery and one hold-out sample, the PGS *R*^2^ is quadratically proportional to the CGR.

**Figure 1.**
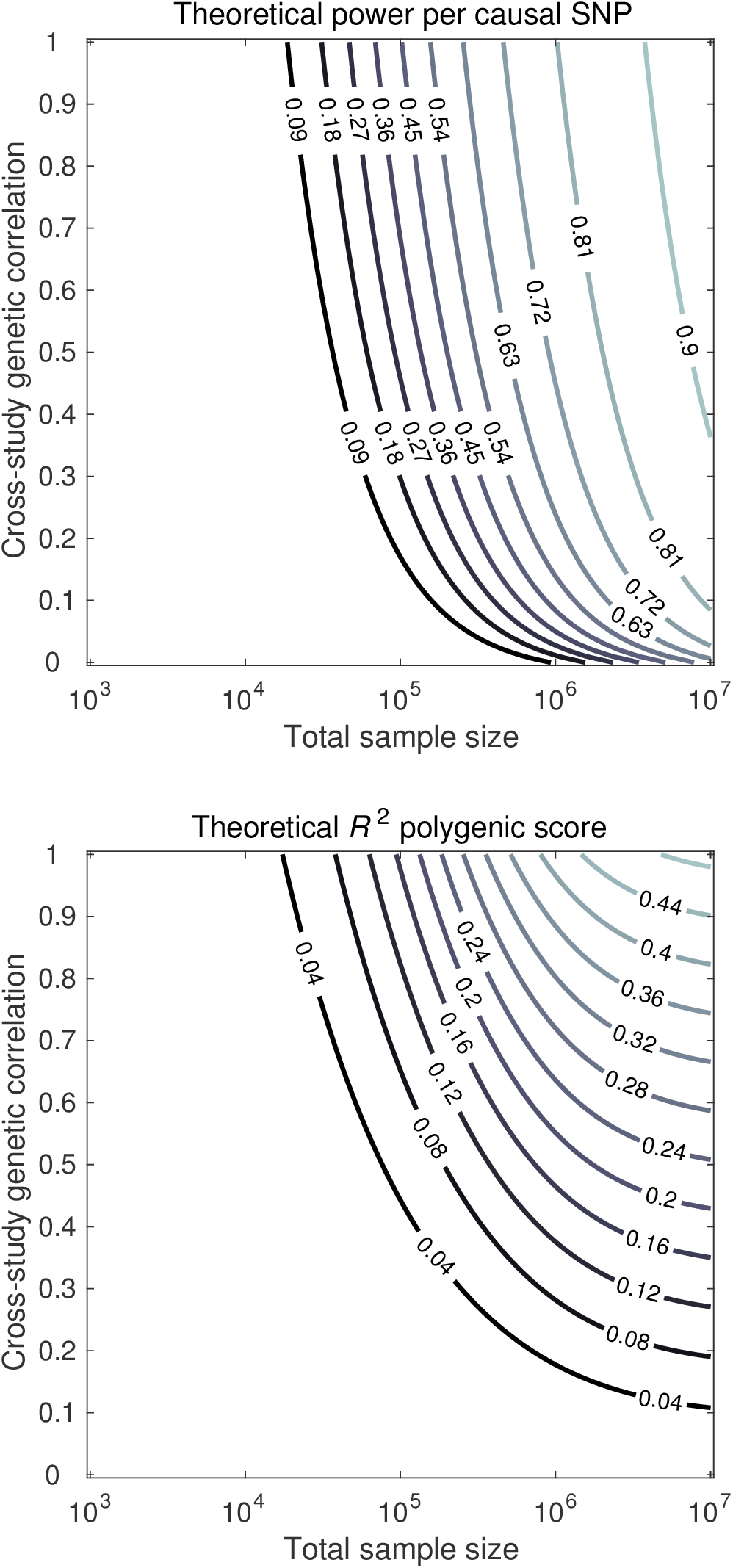
Theoretical predictions of power per causal SNP (upper panel) and out-of-sample *R*^2^ of the PGS (lower panel), for total sample size (*x*-axis) and cross-study genetic correlation (*y*-axis). Factor levels: 50 studies, 100k independent SNPs, and 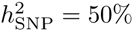 arising from a subset of 1k independent SNPs.

#### SNP heritability and CGR

Fig. 2 shows contour plots for the power per truly associated SNP and *R*^2^ for a setting with 50 studies, with a total sample size of 250,000 individuals, for 1k causal SNPs and 100k SNPs in total, for various combinations of 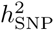 and CGR. The figure shows a symmetric response of both power and *R*^2^ to CGR and 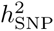. For instance, when 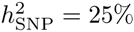 and CGR=0.5 across all studies, the power is expected to be around 34% and the *R*^2^ 3.0%. When these numbers are interchanged (i.e., 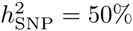 and CGR=0.25), similarly, the power is expected to be 35% and the *R*^2^ 2.9%. Hence, in terms of both *R*^2^ and power, a low heritability can be compensated by a high CGR (e.g., by means of homogeneous measures across studies) and a low CGR can be compensated by high heritability. When either CGR or heritability is equal to zero, both power and *R*^2^ are decimated in the multi-study setting. However, when both are moderately low but still substantially greater than zero, neither power nor *R*^2^ are completely diminished.

**Figure 2.**
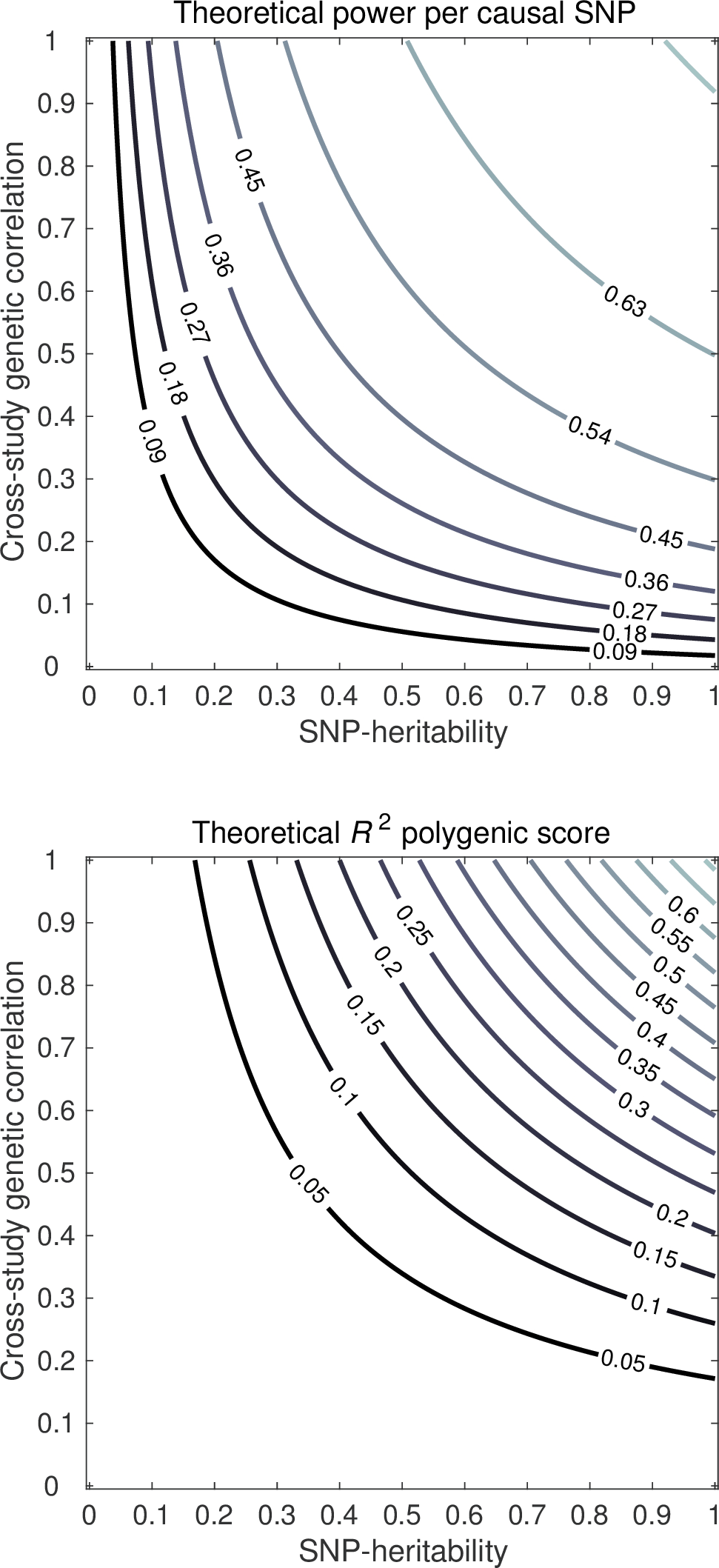
Theoretical predictions of power per causal SNP (upper panel) and out-of-sample *R*^2^ of the PGS (lower panel), for a trait that across studies has SNP heritability (*x*-axis) and cross-study genetic correlation (*y*-axis). Factor levels: 50 studies, sample size 5,000 individuals per study, 100k independent SNPs, and heritability arising from a subset of 1k independent SNPs.

#### Number of studies and CGR

Fig. 3 shows contour plots for the power per truly associated SNP and *R*^2^ for a trait with 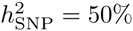, 1k causal SNPs, 100k SNPs in total, and a fixed total sample size of 250,000 individuals. In this figure, various combinations of the CGR and the number of studies are considered. Logically, when there is just one study for discovery, CGR does not affect power. However, even for two studies, the effect of CGR on power is quite pronounced. For instance, when CGR is a half, the power per causal SNP is 63% for one study, 58% for two studies, 51% for ten studies, and 50% for 100 studies. Thus, when the number of studies is low, increasing the number of studies makes the effect of CGR on power more pronounced rapidly. When the number of studies is large, further increases in the number of studies hardly make the effect of CGR on power more pronounced.

**Figure 3.**
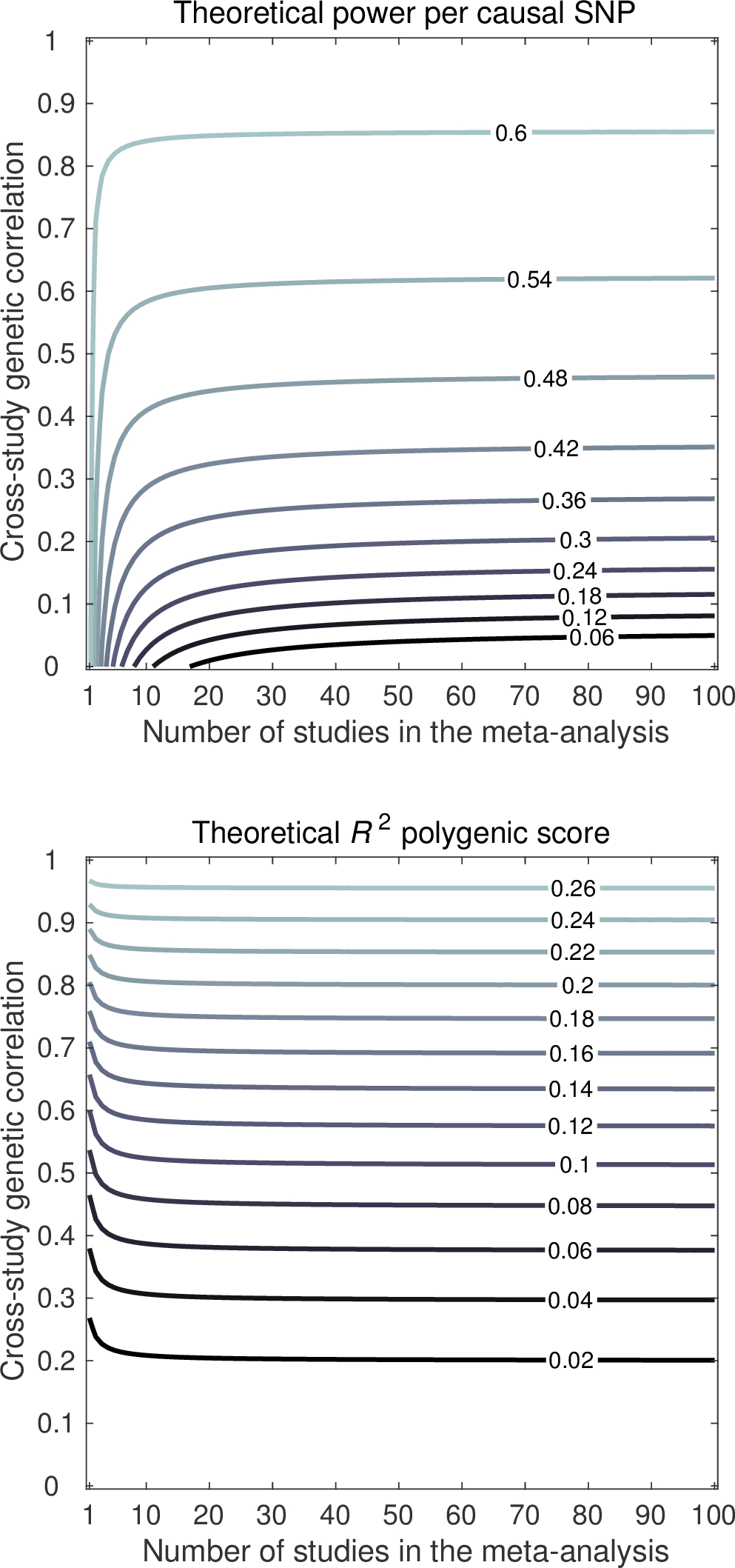
Theoretical predictions of power per causal SNP (upper panel) and out-of-sample *R*^2^ of the PGS (lower panel), for a trait with GWAS results from the number of studies (*x*-axis) with cross-study genetic correlation (*y*-axis). Factor levels: total sample size 250,000 individuals, 100k independent SNPs, and heritability 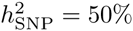 arising from a subset of 1k independent SNPs.

For a given number of studies, we observed that the effect CGR has on *R*^2^ is stronger than the effect it has on power. This observation is in line with the approximated theoretical *R*^2^ in Eq. 2, indicating that *R*^2^ is quadratically proportional to CGR. However, an interesting observation is that this quadratic relation lessens as the number of studies grows large, despite the total sample size being fixed. For instance, at a CGR of a half, the *R*^2^ in the hold-out sample is expected to be 6.9% when there is only one discovery study. However, the expected *R*^2^ is 8.1% for two discovery studies, 9.3% for ten discovery studies, and 9.6% for 100 discovery studies. The reason for this pattern is that, in case of one discovery study, the PGS is influenced relatively strongly by the study-specific component of the genetic effects. This idiosyncrasy is not of relevance for the hold-out sample. As the number of studies increases – even though each study brings its own idiosyncratic contribution – each study consistently conveys information about the part of the genetic architecture which is common across the studies. Since the idiosyncratic contributions from the studies are independent, they tend to average each other out, whereas the common underlying architecture gets more pronounced as the number of studies in the discovery increases, even if the total sample size is fixed.

#### SNP heritability in the hold-out sample

Fig. 4 shows a contour plot for the PGS *R*^2^ based on a meta-analysis of 50 studies with a total sample size of 250,000 individuals, with 1k causal SNPs and 100k SNPs in total, and a CGR of 0.8 between both the discovery studies and the hold-out sample. In the plot, various combinations of 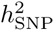 in the discovery samples and 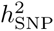 in the hold-out sample are considered. The response of PGS *R*^2^ to heritability in the discovery sample and the hold-out sample is quite symmetric, in the sense that a low 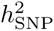 in the discovery samples and a high 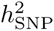 in the hold-out sample yield a similar *R*^2^ as a high 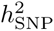 in the discovery sample and a low 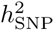 in the hold-out sample. However, *R*^2^ is slightly more sensitive to 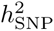 in the hold-out sample than in the discovery samples. For instance, when SNP heritability in the discovery samples is 50% and 25% in the hold-out sample, the expected *R*^2^ is 10%, whereas in case the SNP heritability is 25% in the discovery samples and 50% in the hold-out sample, the expected *R*^2^ is 13%.

**Figure 4.**
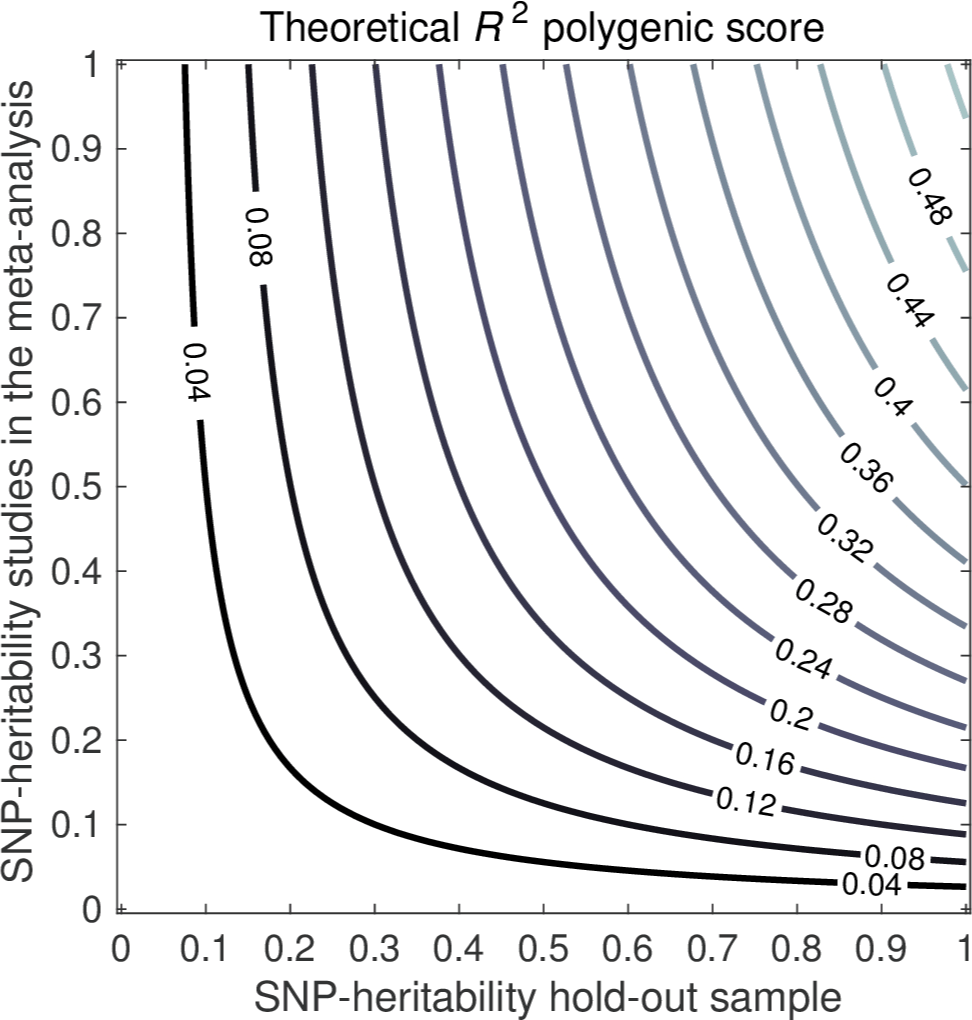
Theoretical predictions of out-of-sample *R*^2^ of the PGS, for the SNP heritability in the hold-out sample (*x*-axis) and the SNP heritability in the discovery samples (*y*-axis). Factor levels: 50 studies, sample size 5,000 individuals per study, cross-study genetic correlation 0.8, 100k independent SNPs, and heritability arising from a subset of 1k independent SNPs.

#### CGR between sets of studies

Fig. 5 shows a contour plot for the power per truly associated SNP in a setting where there are two sets consisting of 50 studies each. Within each set, the CGR is equal to one, whereas between sets the CGR is imperfect. Consider, for example, a scenario where one wants to meta-analyze GWAS results for height from a combination of two sets of studies; one set of studies consisting primarily of individuals of European ancestry and one set of studies with mostly people of Asian ancestry in it. Now, one would expect CGRs close to one between studies consisting primarily of individuals of European ancestry and the same for the CGRs between studies consisting primarily of people of Asian ancestry. However, the CGRs between those two sets of studies may be less than one.

**Figure 5.**
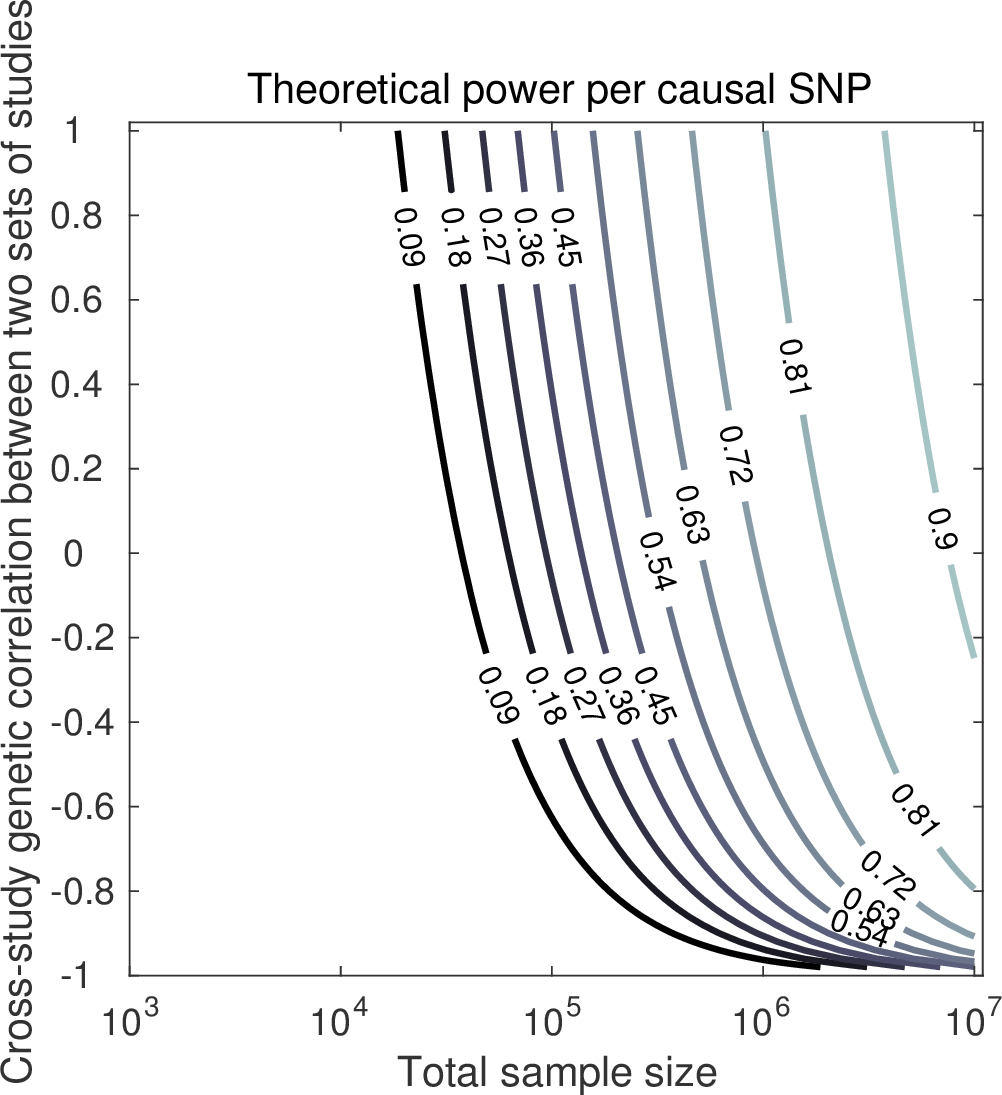
Theoretical predictions of power per causal SNP, for total sample size (*x*-axis) and CGR between two sets of studies (*y*-axis). Factor levels: 2 sets of 50 studies, CGR equal to 1 within both sets, 100k independent SNPs, and heritability 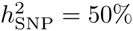 arising from a subset of 1k independent SNPs.

As is shown in S1 Derivations Power, in case the CGR between the two sets of studies, 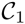 and 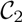, is zero, meta-analyzing the two sets jointly yields power 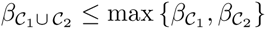 and 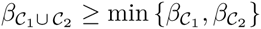, where 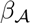 denotes the power in set of studies 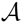. In particular, when 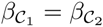 we have under a CGR of zero between the sets, that 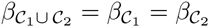. Since in Fig. 5 we considered two equally-powered sets, the power of a meta-analysis using both sets, under zero CGR between sets, is identical to the power obtained when meta-analyzing, for instance, only the first set. However, as CGR between sets increases, so does power. For instance, when a total sample size of 250,000 individuals is spread across 2 clusters, each cluster consisting of 50 studies (i.e., sample size of 125,000 individuals per cluster and 2,500 individuals per study), under 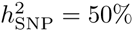 due to 1k causal SNPs, a CGR of one within each cluster, and CGR of zero between clusters, the power is expected to be 49%, which is identical to the power of a meta-analysis of either the first or the second cluster. However, if the CGR between clusters is 0.5 instead of zero, the power goes up to 58%. In terms of the expected number of hits, this cross-ancestry meta-analysis yields an expected 82 additional hits, compared to a meta-analysis considering only one ancestry.

Alternatively, one could carry out a meta-analysis in each set of studies and pool the hits across these sets. However, this would imply more independent tests being carried out, and, hence, the need for a more stringent genome-wide significance threshold, in order to keep the false-postive rate fixed. Therefore, this route may yield less statistical power than a meta-analysis of merely one of the two sets or a joint analysis of both. Ideally, in the scenario where between-population heterogeneity is likely, one should apply a meta-analysis method that accounts for the heterogeneity (e.g., [27–31]). By applying such a method, one can consider all GWAS results from different ancestry groups in one analysis.

## Empirical results for SNP-based heritability and CGR

In Table 1 we report univariate GREML estimates of SNP heritability and bivariate GREML estimates of genetic correlation for traits that attained a pooled sample size of at least 18,000 individuals, which gave us at least 50% power to detect a genetic correlation near one for a trait that has a SNP heritability of 10% or more [43]. The smallest sample size is *N*=19,184 for self-rated health. Details per phenotype (i.e., sample size, univariate estimates of SNP heritability, and bivariate estimates of genetic correlation, stratified across studies and sexes, as well as cross-study and cross-sex averages) are provided in S7 GREML Results.

**Table 1.**
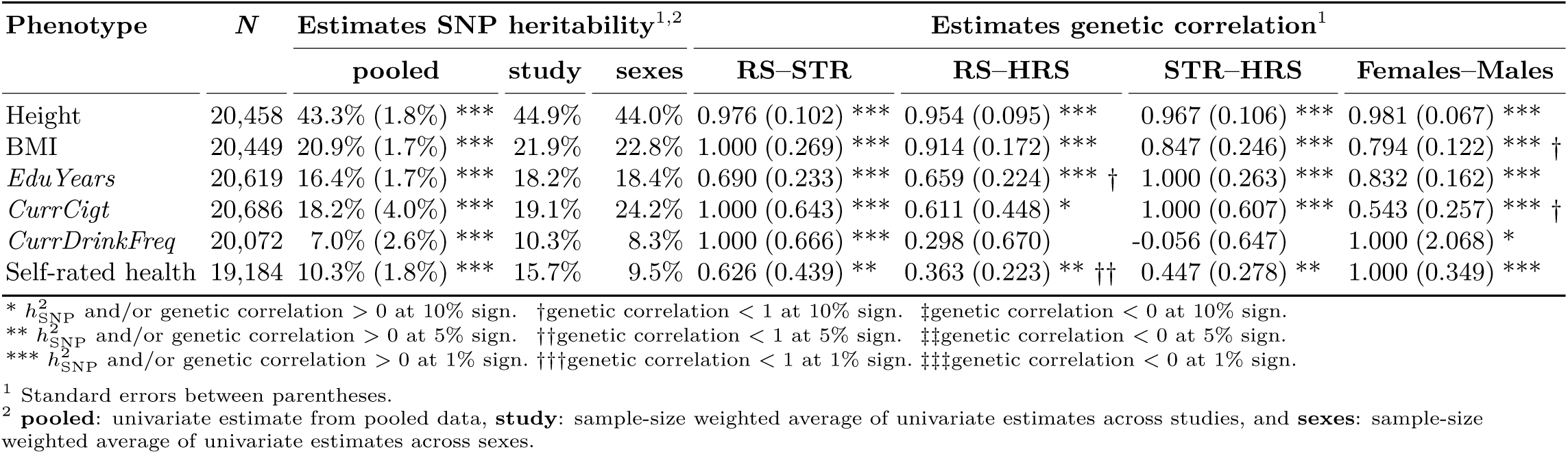
GREML estimates of SNP heritability 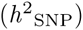 and genetic correlation across studies and sexes.

The univariate estimates of SNP heritability based on the pooled data assume perfect CGRs. Therefore, such estimates of SNP heritability are downwards biased when based on data from multiple studies with imperfect CGRs. To circumvent this bias, we estimated SNP heritability in each study separately, and focused on the sample-size-weighted cross-study average estimate of SNP heritability.

For both height and BMI, we observed genetic correlations close to one across pairs of studies and between females and males. For years of schooling (*EduYears*) we found a CGR around 0.8 when averaged across pairs of studies. Similarly, the genetic correlation for *EduYears* in females and males lies around 0.8. The CGR of self-rated health is substantially below one across the pairs of studies, whilst the genetic correlation between females and males seems to lie around one. The reason for this difference in the genetic correlation of self-rated health between pairs of studies and between females and males may be due to the difference in the questionnaire across studies, discussed in S5 Data and Quality Control. The questionnaire differences can yield a low CGR, while not precluding the remaining genetic overlap for this measure across the three studies, to be highly similar for females and males. For *CurrCigt* and *CurrDrinkFreq*, the estimates of CGR and of genetic correlation between females and males are non-informative. For these two traits the standard errors of the genetic correlations estimates are large, mostly greater than 0.5. In addition, for *CurrDrinkFreq* there is strong volatility in the CGR estimate across pairs of studies.

## Attenuation in power and *R*^2^ due to imperfect CGR

Considering only the traits for which we obtained accurate estimates of CGR and SNP heritability (i.e., with low standard errors), we used the MetaGAP calculator to predict the number of hits in a set of discovery samples and the PGS *R*^2^ in a hold-out sample, in prominent GWAS efforts for these traits. Details and notes on the results from existing studies, used as input for the MetaGAP calculations, can be found in S8 Large-scale GWAS efforts. Importantly, for the traits under consideration here, all large-scale GWAS results obtained using a meta-analysis, use a fixed-effects meta-analysis.

Since we only had accurate estimates for height, BMI, *EduYears*, and self-rated health, we focused on these four phenotypes. For these traits, we computed sample-size-weighted average CGR estimates across the pairs of studies. Table 2 shows the number of hits and PGS *R*^2^ reported in the most comprehensive GWAS efforts to date for the traits of interest, together with predictions from the MetaGAP calculator. We tried several values for the number of independent haplotype blocks (i.e., 100k, 150k, 200k, 250k) and for the number of trait-associated blocks (i.e., 10k, 15k, 20k, 25k). Overall, 250k blocks of which 20k trait-affecting yielded theoretical predictions in best agreement with the empirical observations; we acknowledge the potential for some overfitting (i.e., two free parameters set on the basis of 17 data points; 10 data points for the reported number of hits and 7 for PGS *R*^2^).

**Table 2.**
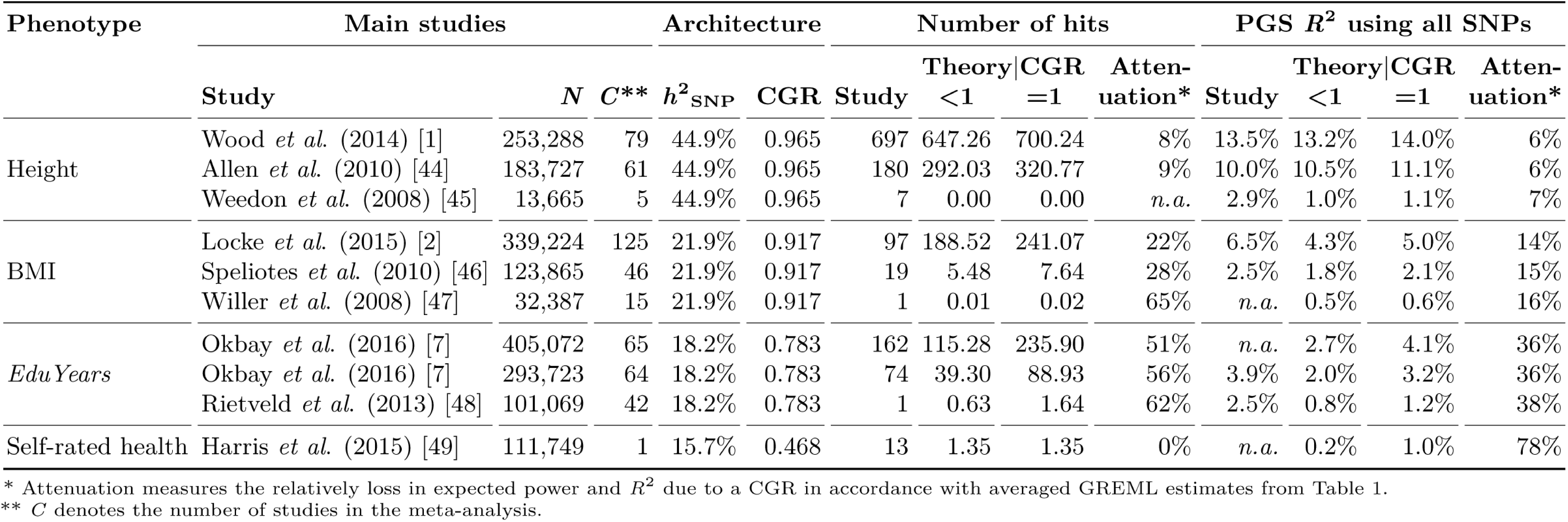
Predicted and observed number of genome-wide-significant hits and PGS *R*^2^, for large-scale GWAS efforts to date for height, BMI, *EduYears*, and self-rated health, assuming 250k effective SNPs (i.e., independent haplotype blocks) of which 20k trait-affecting, using averaged GREML estimates from Table 1 for setting SNP heritability and CGR. Notes on the sources for the large-scale GWAS efforts are listed in Table 10.

For height – the trait with the lowest standard error in the estimates of 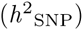 and CGR – the predictions of the number of hits and PGS *R*^2^ for the two largest GWAS efforts are much in line with theoretical predictions. For the smaller GWAS of 13,665 individuals [45], our estimates seem slightly conservative; 0 hits expected versus the 7 reported. However, in our framework, we assumed that each causal SNP has the same *R*^2^. Provided there are some differences in *R*^2^ between causal SNPs, the first SNPs that are likely to reach genome-wide significance in relatively small samples, are the ones with a comparatively large *R*^2^. This view is supported by the fact that a PGS based on merely 20 SNPs already explains 2.9% of the variation in height. Hence, for relatively small samples our theoretical predictions of power and *R*^2^ may be somewhat conservative. In addition, the 10k SNPs with the lowest meta-analysis *p*-values can explain about 60% of the SNP heritability [1]. If the SNPs tagging the remaining 40% each have similar predictive power as the SNPs tagging the first 60%, then the number of SNPs needed to capture the full 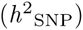 would lie around 10k/0.6=17k, which is somewhat lower than the 20k which yields the most accurate theoretical predictions. However, as indicated before, the SNPs which appear most prominent in a GWAS are likely to be the ones with a greater than average predictive power. Therefore, the remaining 40% of 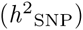 is likely to be stemming for SNPs with somewhat lower predictive power. Hence, 20k associated independent SNPs is not an unlikely number for height.

The notion of a GWAS first picking up the SNPs with a relatively high *R*^2^ is also supported by the predicted and observed number of hits for the reported self-rated-health GWAS [49]; given a SNP heritability estimate between 10% [49] and 16% (Table 2), according to our theoretical predictions, a GWAS in a sample of around 110k individuals is unlikely to yield even a single genome-wide significant hit. Nevertheless, this GWAS has yielded 13 independent hits. This finding supports the idea that for various traits, some SNPs with a relatively high *R*^2^ are present. However, there is uncertainty in the number of truly associated loci. More accurate estimates of this number may improve the accuracy of our theoretical predictions.

For BMI our predictions of PGS *R*^2^ were quite in line with empirical results. However, for the number of hits, our predictions for the largest efforts seemed overly optimistic. We therefore suspect that the number of independent SNPs associated with BMI is higher than 20k; as a higher number of associated SNPs would reduce the GWAS power, while preserving PGS *R*^2^, yielding good agreement with empirical observation. Nevertheless, given the limited number of data points, this strategy of setting the number of causal SNPs would increase the chance of overfitting.

For *EduYears* we observed that the reported number of hits is in between the expected number of hits when the CGR is set to the averaged GREML estimate of 0.783 and when the CGR is set to one. Given the standard errors in the CGR estimates for *EduYears*, the CGR might very well be somewhat greater than 0.783, which would yield a good fit with the reported number of hits. However, as with the number of truly associated SNPs for BMI, in light of the risk of overfitting, we can make no strong claims about a slightly higher CGR of *EduYears*.

Overall, our theoretical predictions of the number of hits and PGS *R*^2^ are in moderate agreement with empirical observations, especially when bearing in mind that we are looking at a limited number of data points, making chance perturbations from expectation likely. In addition, regarding the number of hits, the listed studies are not identical in terms of the procedure to obtain the independent hits. Therefore, the numbers could have been slightly different, had the same pruning procedure been used across all reported studies.

Regarding attenuation, we observed a substantial spread in the predicted number of hits and PGS *R*^2^ when assuming either a CGR equal to one, or a CGR in accordance with empirical estimates, with traits with lower CGR suffering from stronger attenuation in power and predictive accuracy. In line with theory, *R*^2^ falls sharply with CGR. For instance, for self-rated health, the estimate CGR of about 0.5, would – in expectation – yield a PGS that retains only 0.5^2^=25% of the *R*^2^ it would have had under a CGR of one. This is supported by the reported attenuation of roughly 80%.

Given our CGR estimates, we expect a relative loss in PGS *R*^2^ of 6% for height, 14% for BMI, 36% for *EduYears*, and 78% for self-rated health, compared to the *R*^2^ of a PGS under perfect CGRs (Table 2). This loss in *R*^2^ is unlikely to be reduced by larger sample sizes and denser genotyping.

Somewhat contrary to expectation, the number of hits seems to respond even more strongly to CGR than PGS *R*^2^. However, since in each study under consideration the average power per associated SNP is quite small, a small decrease in power per SNP in absolute terms can constitute a substantial decrease in relative terms. For instance, when one has 2% power per truly associated SNP, an absolute decrease of 1% – leaving 1% power – constitutes a relative decrease of 50% of power per causal SNP, and thereby a 50% decrease in the expected number of hits. This strong response shows, for example, in the case of *EduYears*, where the expected number of hits drop by about 37% when going from a CGR of one down to a CGR of 0.783.

## Discussion

We aimed to answer the question whether imperfect cross-study genetic correlations (CGRs) help to explain a part of the ‘hiding heritability’ for traits such as height. We showed that imperfect CGRs are indeed likely to contribute to the gap between the phenotypic variation accounted for by all SNPs jointly and by the leading GWAS efforts to date. We arrived at this conclusion in five steps.

First, we developed a Meta-GWAS Accuracy and Power (MetaGAP) calculator that accounts for the CGR. This online calculator relates the statistical power to detect associated SNPs and the *R*^2^ of the polygenic score (PGS) in a hold-out sample to the number of studies, sample size and SNP heritability per study, and the CGR. The underlying theory shows that there is a quadratic response of the PGS *R*^2^ to CGR. Moreover, we showed that the power per associated SNP is also affected by CGR.

Second, we used simulations to demonstrate that our theory is robust to several violations of the assumptions about the underlying data-generating process, regarding the relation between allele frequency and effect size, the distribution of allele frequencies, and the factors contributing to CGR. Further research needs to assess whether our theoretical predictions are also accurate under an even broader set of scenarios (e.g., when studying a binary trait).

Third, we used a sample of unrelated individuals from the Rotterdam Study, the Swedish Twin Registry, and the Health and Retirement Study, to estimate SNP-based heritability as well as the CGR for traits such as height and BMI. Although our CGR estimates have considerable standard errors, the estimates make it likely that for many polygenic traits the CGR is positive, albeit smaller than one.

Fourth, based on these empirical estimates of SNP heritability and CGR for height, BMI, years of education, and self-rated health, we used the MetaGAP calculator to predict the number of expected hits and the expected PGS *R*^2^ for the most prominent studies to date for these traits. We found that our predictions are in moderate agreement with empirical observations. Our theory seems slightly conservative for smaller GWAS samples. For large-scale GWAS efforts our predictions were in line with the outcomes of these efforts. More accurate estimates of the number of truly associated loci may further improve the accuracy of our theoretical predictions.

Fifth, we used our theoretical model to assess statistical power and predictive accuracy for these GWAS efforts, had the CGR been equal to one for the traits under consideration. Our estimates of power and predictive accuracy in this scenario indicated a strong decrease in the PGS *R*^2^ and the expected number of hits, due to imperfect CGRs. Though these observations are in line with expectation for predictive accuracy, for statistical power the effect was larger than we anticipated. This finding can be explained, however, by the fact that though the absolute decrease in power per SNP is small, the relative decrease is large, since the statistical power per associated SNP is often low to begin with.

Overall, our study affirms that although PGS accuracy improves substantially with further increasing sample sizes, in the end PGS *R*^2^ will continue to fall short of the full SNP-based heritability. Hence, this study contributes to the understanding of the hiding heritability reported in the GWAS literature.

Regarding the etiology of imperfect CGRs, the likely reasons are heterogeneous phenotype measures across studies, gene–environment interactions with underlying environmental factors differing across studies, and gene–gene interactions where the average effects differ across studies due to differences in allele frequencies. Our study is not able to disentangle these different causes; by estimating the CGR for different traits we merely quantify the joint effect these three candidates have on the respective traits.

However, in certain situations it may be possible to disentangle the etiology of imperfect CGRs to some extent. For instance, in case one considers a specific phenotype that is usually studied by means of a commonly available but relatively heterogeneous and/or noisy measure, while there also exists a less readily available but more accurate and homogeneous measure. If one has access to both these measures in several studies, one can compare the CGR estimates for the more accurate measure and the CGR estimates for the less accurate but more commonly available measure. Such a comparison would help to disentangle the contribution of phenotypic heterogeneity and genetic heterogeneity to the CGR of the more commonly available measure.

In considering how to properly address imperfect CGRs, it is important to note that having a small set of large studies, rather than a large set of small studies, does not necessarily abate the problem of imperfect genetic correlations. Despite the fact that having fewer studies can help to reduce the effects of heterogeneous phenotype measures, larger studies are more likely to sample individuals from different environments. If gene–environment interactions do play a role, strong differences in environment between subsets of individuals in a study can lead to imperfect genetic correlations within that study. The attenuation in power and accuracy resulting from such within-study heterogeneity may be harder to address than cross-study heterogeneity.

Our findings stress the importance of considering the use more sophisticated meta-analysis methods that account for cross-study heterogeneity [27–31]. We believe that the online MetaGAP calculator will prove to be an important tool for assessing whether an intended fixed-effects meta-analysis of GWAS results from different studies is likely to yield meaningful outcomes.

## Supporting Information

### S1 Derivations Power

In this section, we derive an expression for the power of a meta-analysis of GWAS results, under a design with many studies, with arbitrary sample sizes, SNP-based heritability, and cross-study genetic correlation (CGR).

First, the underlying assumptions are presented. Second, we write the GWAS *Z* statistics in terms of the true SNP effect and noise. Third, we incorporate cross-study genetic correlations by assuming a model with random SNP effects that are correlated imperfectly across studies. Using the Cholesky decomposition of the cross-study genetic correlation matrix, we write the correlated SNP effects in terms of a weighted sum of independent genetic factors. By means of this decomposition into independent factors, we derive the distribution of the *Z* statistic in a given study, as well as the distribution of the multi-study meta-analysis *Z* statistic. From the latter distribution we obtain a framework for performing multi-study power calculations.

It is important to note that models which incorporate random SNP effects have been widely used, for instance, to estimate variance components [35] and genetic correlations across traits and samples [42], to control for cryptic relatedness and population structure in a GWAS [41], and to distill the constituents of genomic inflation [36, 40]. Hence, the novelty in our work lies not in using random SNP-effect models to incorporate imperfect genetic correlations across studies. Instead the novelty lies in the subsequent step, viz., to use such models in order to perform power calculations under the presence of imperfect CGRs.

#### Assumptions

We derive an expression of statistical power for a quantitative trait in sample-size weighted meta-analysis [34]. In order to arrive at a tractable expression of statistical power, we make the following assumptions.

1. When considering a given SNP in the GWAS, any phenotypic variance due to other SNPs gets absorbed by the normally, independent, and identically distributed residual term (which is what happens when studying a sample of unrelated individuals, and which is in line with assumptions underlying most GWAS packages, except for family-based and mixed-linear-model-type GWAS software). This assumption keeps the algebra simple at the cost of a small loss in generality. In S4 Simulation Studies we show that violations of this assumption do not affect results.
2. The regressors (i.e., SNP data) in the meta-analysis studies are fixed (i.e., non-stochastic)—this assumption is equivalent to conditioning on the genotype data. This assumption also keeps the algebra simple at the cost of a small loss in generality. In S4 Simulation Studies we show that violations of this assumption do not affect results.
3. Each causal locus is shared across all studies. This assumption enables us to consider CGRs as a one-dimensional factor that is shaped solely by the cross-study correlation of the effects of trait-affecting haplotype blocks. In S4 Simulation Studies we show that violations of this assumption hardly affect results.
4. The genome can be divided into independent haplotype blocks, where for each block we have precisely one SNP that tags all the variation within this block. By means of this assumption, we can ignore linkage disequilibrium, thereby strongly reducing the complexity of our derivations. In addition, we assume that the effects of trait-affecting haplotype blocks are independent. The former assumption would imply that all trait-affecting variation in a haplotype block can be captured by the single tag SNP for that block. Although we make no claim that common SNPs perfectly tag all trait affecting variants, we do claim that a relatively small set of common SNPs can tag the heritability as estimated using common SNPs. Consequently, when using estimates of SNP heritability based on common SNPs, we deem this assumption and its implications to generate little bias in our theoretical predictions.
5. The effect sizes of SNPs are inversely related to SNP variance (i.e., rare variants have larger effects than common variants, such that the expected *R*^2^ of each causal SNP, with respect to the phenotype, is equal regardless of allele frequency). This assumption makes it possible to compute statistical power without having to specify the allele frequency and an *a priori* unknown effect size. Under this assumption, SNP heritability and the number of trait-affecting haplotype blocks replace a SNP-specific effect size and allele frequency. In S4 Simulation Studies we show that violations of this assumption hardly affect results.

#### Single-SNP model

Here, we write the GWAS *Z* statistic in a given study for a given SNP, as a function of the true effect and noise. This decomposition into true effect and noise helps to derive the distribution of the *Z* statistic.

For studies *j*=1,…,*C* and SNPs *k*=1,…,*S*, let the model for a quantitative trait with a single SNP as predictor (Assumption 1) for the mean-centered phenotype y_*j*_ be given by

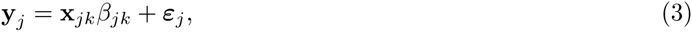

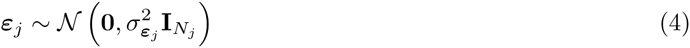

where *X*_*jk*_ denotes the mean-centered genotype vector of SNP *k* in study *j*, scaled such that 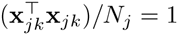. In Eq. 3, *β*_*jk*_ is the effect of SNP *k* in study *j*. In Eq. 4, *ε*_*j*_ is the residual and 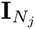 the *N*_*j*_ × *N*_*j*_ identity matrix, where *N*_*j*_ denotes the sample size of study *j*.

The GWAS estimate of *β*_*jk*_ for a quantitative trait is usually obtained by applying OLS. Hence, it can be written as

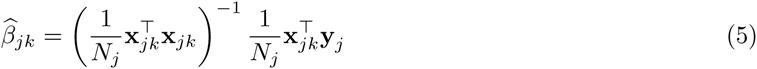

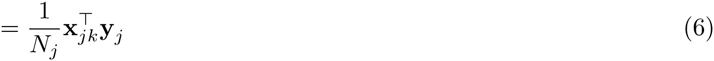

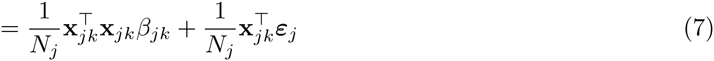

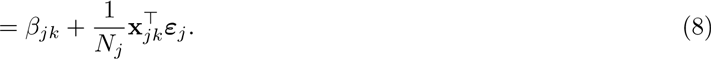

Using standard results from regression theory assuming fixed regressors (Assumption 2) and the aforementioned scaling of the genotype vector, the theoretical variance of the OLS-estimate of the SNP effect is given by

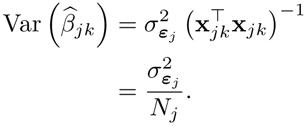

Therefore, the standard error of the OLS estimate is given by

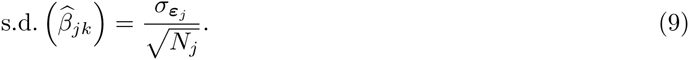

By taking the ratio of Eq. 8 and 9 we obtain the *Z* statistic (instead of the commonly used and highly similar *t*-test statistics) for SNP *k* in study *j*. That is,

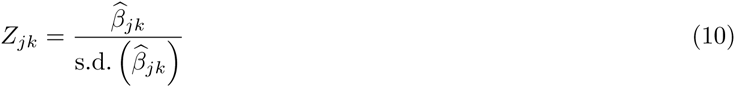

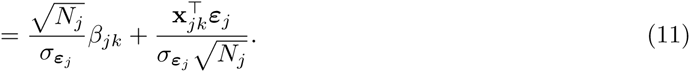

Let *v*_*jk*_ denote the last term in the right-hand side of Eq. 11. Under the aforementioned scaling of the regressor and the distribution of *ε*_*j*_, it follows from standard properties of the multivariate normal distribution that 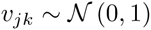.

#### Modelling cross-study genetic correlation

We incorporate cross-study genetic correlations by considering a model with random SNP effects, correlated across studies. For ease of derivations, we assume that each causal SNP contributes across all studies (Assumption 3). In order to simplify further derivations, we use a Cholesky decomposition to write correlated SNP effects in terms of independent underlying factors. Using this independent-factor representation, we derive the distribution of a GWAS *Z* statistic, in terms of the study-specific noise and contributions of the underlying genetic factors.

Genetic correlation can be conceptualized as the correlation between SNP effects across different strata (e.g., across populations, studies, age groups, etc.). Taking studies as ‘strata’, a group of *C* studies has *C* × *C* genetic correlation matrix, denoted by ***P***_G_.

When effects are normally distributed, a given correlation structure between effects is most straightforwardly obtained by constructing the Cholesky decomposition of the correlation matrix, and multiplying independent standard-normal random variables by this decomposition. An interpretation of this decomposition is that it provides a set of weights that transform a set of independent underlying genetic factors into correlated genetic effects.

First, we formalize how to transform independent standard-normal random variables into correlated normal random variables. Let **Γ**_G_ be the lower-triangular Cholesky decomposition of the genetic correlation matrix, such that 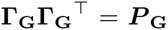, let 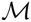 denote the set of *M* causal SNPs, let **E** be an *C*-by-*M* matrix of independent standard normal draws from different genetic factors (rows) for the different causal SNPs (columns), and let *η*_*k*_ be the column of **E** corresponding to causal SNP *k*. Then

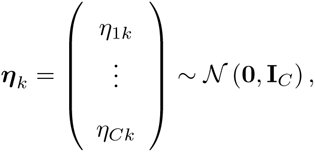

where *η*_*k*_ is independent of *η*_*l*_ for *l* ≠ *k* (Assumption 4). Now, for SNP *k* in the set of causal SNPs, we can define the vector of effects across studies for the given SNP, such that it has correlation matrix ***P***_G_, as follows:

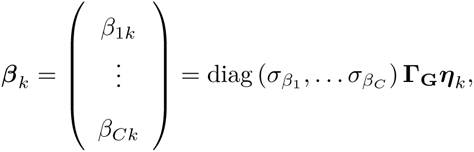

where diag() is a diagonal matrix with specified elements as diagonal entries, and

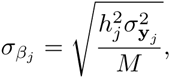

with 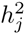 (resp. 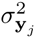) denoting the SNP heritability (phenotypic variance) in study *j*. Under this design of study-specific SNP effects, we attain a CGR structure in line with *P*_G_ and the desired study-specific SNP heritabilities.

Using this approach for constructing correlated SNP effects, we can write the effect of SNP *k* in study *j* (i.e., *β*_*jk*_) as a linear combination of the independent underlying 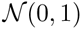 distributed random variables. That is,

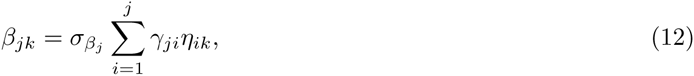

where γ*_ji_* denotes element in row *j* column *i* of **Γ** and *η*_*ik*_ the *i*-th element of *η*_*k*_. Given our scaling of SNPs, the *R*^2^ of each causal SNP in study *j* is given by 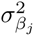, regardless of the allele frequency of the SNP of interest (Assumption 5).

We can now write the GWAS *Z* statistic for a given SNP in a given study, as a linear combination of independent random variables. For SNP *k* in the set of *P* non-causal SNPs, denoted by 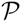 (such that 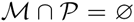), we have for all studies *j* that *β*_*jk*_=0. By substituting *β* in Eq. 11 according to Eq. 12 for causal SNPs and the preceding equality for non-causal SNPs, we obtain the following expression for the *Z* statistic of SNP *k* in study *j*:

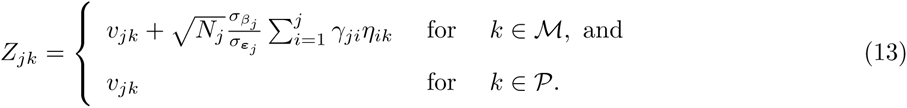

#### Distribution meta-analysis *Z* statistic

Here, we derive the distribution of the meta-analysis *Z* statistic and reduce the number of input parameters by appropriate substitutions. Finally, for intuition, we present the distribution of *Z* statistics from a meta-analysis of GWAS results from two studies.

For any SNP *k* in the set 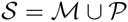 of *S*=*M* + *P* causal and non-causal SNPs, we use the sample-size-weighted meta-analysis *Z* statistic [34], defined as follows:

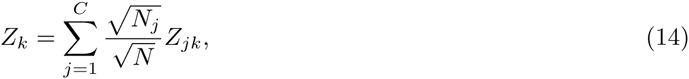

where *N*=*N*_1_ + … + *N*_*C*_ denotes the total sample size. Plugging Eq. 13 for 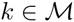 into Eq. 14, yields an expression for the meta-analysis *Z* statistic in terms of independent random variables. That is,

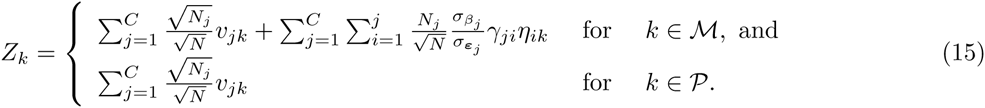

As the *υ*_*jk*_ terms in the preceding expression are independent standard-normal draws, it follows that

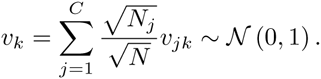

In Eq. 15 we have a double sum over random variables. However, by changing the order of summation, this double sum can be rewritten as follows:

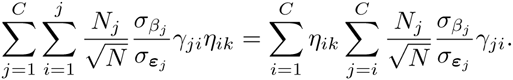

Therefore, we can rewrite Eq. 15 as follows:

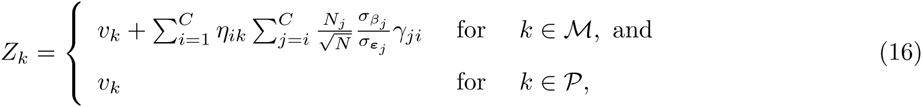

where the inner sum yields the weight for the random variable of interest.

Exploiting the fact that *η*_*ik*_ and *v*_*k*_ are independent standard-normal draws, the variance of the sum of terms is equal to the sum of the variance of the respective terms. Hence, we have that

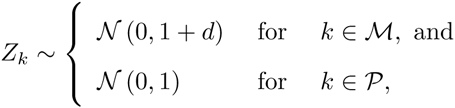

where

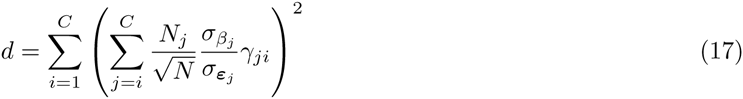

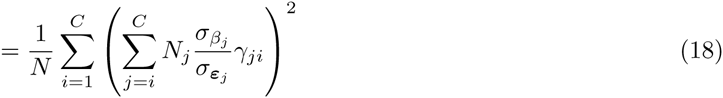

The quantity *d* we refer to as the ‘power parameter’. Since this parameter is a sum of squares, it is non-negative. The greater the power parameter is, the higher statistical power the meta-analysis of GWAS results has. Note that in case 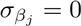 for all *j* (i.e., the trait is not heritable in any study), that *d*=0, and hence the meta-analysis *Z* statistic reverts to a standard-normal test statistic, which matches the distribution under the null. However, as 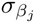 increases, *d* becomes larger, yielding a meta-analysis with higher statistical power.

Given SNP-based heritability, phenotypic variation, and the number of causal variants, we have that the effect size per causal SNP in a study is given by 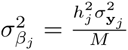, and the residual variance, absorbing the variance due to the omitted *M* – 1 SNPs (Assumption 1), is given by 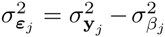. Using these expressions, we can write the ratio of 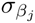 and 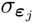, appearing in Eq. 18, as a function of only heritability and the number of causal SNPs. That is,

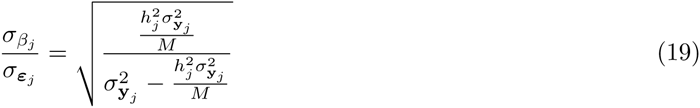

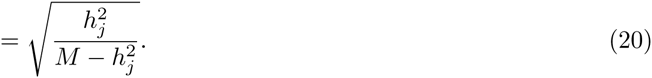

Plugging the last expression into Eq. 18 yields

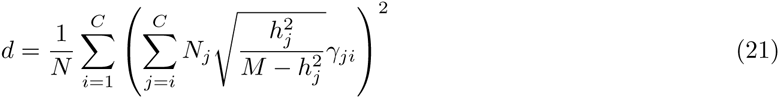

This expression for the power parameter shows that it is not affected by scaling due to phenotypic variance; the parameter is only affected by the cross-study genetic correlation matrix, the SNP-based heritability per study, and the sample size per study.

In case the number of studies is two, with sample size *N* in Study 1 and *N* in Study 2, SNP heritability 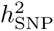, and a genetic correlation *ρ*_G_ between the two studies, we have that the meta-analysis *Z* statistic, of a trait-affecting SNP *k*, is normally distributed with mean zero and

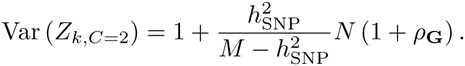

Bearing in mind that the number of causal SNPs *M* ≫ 1 under a highly polygenic model, while *h*^2^ ∈ [0, 1], we have that under high polygenicity 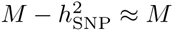. Hence, an easy approximation of the variance of *Z*_*k*_ is given by

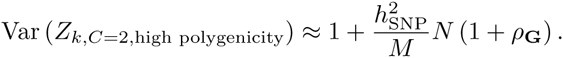

In the scenario where the cross-study genetic correlations equals one, we have that 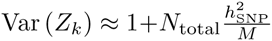 for a trait-affecting haplotype block and Var (*Z*_*k*_)=1 for a non-causal haplotype block, where *N*_total_=2*N*. These expressions are equivalent to the expected value of the squared *Z* statistics from the linear regression analysis reported in Section 4.2 of the Supplementary Note to [41], as well as the first equation in [36] when assuming that confounding biases and linkage disequilibrium are absent.

#### Adding genetically uncorrelated studies to the meta-analysis

Here, we consider what happens to statistical power of a meta-analysis of GWAS results from several sets of studies, with genetic correlations between the studies within each set, but with no genetic correlation between the different sets. We first consider a scenario with one set consisting of *C* – 1 studies and one other set consisting of only one study. We then generalize to a setting with multiple sets, each set containing at least one study. We show that the power parameter for a meta-analysis of several sets of studies with no genetic correlations between sets, can be written as a sample-size weighted sum of the power parameters within the respective sets.

In case one has *C* – 1 studies with associated CGR matrix, the associated Cholesky decomposition denoted by **Γ**_(*C*)_, and an additional study indexed by *C*, which is genetically uncorrelated to the *C* – 1 other studies, then the *C* × *C* Cholesky decomposition of the full CGR matrix is given by

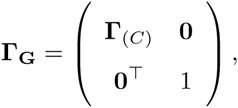

where **0** denotes a column vector of zeros.

Now, the quantity *d* in Eq. 21 can be decomposed as follows.

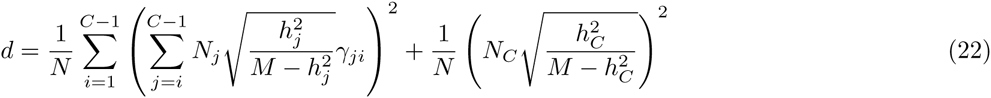

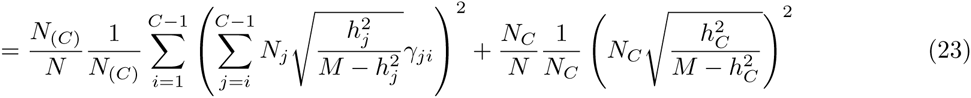

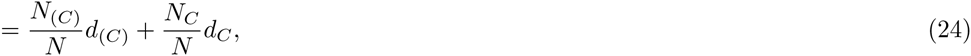

where *d*_*C*_ denotes the power parameter in Eq. 21 had only study *C* (with sample-size *N*_*C*_) be considered, and *d*_(*C*)_ the power parameter in Eq. 21 had only the first *C* − 1 studies (with total corresponding sample-size *N*_(*C*)_) be considered. Hence, the power parameter in this scenario is the sample-size-weighted sum of the power parameter of the first *C* – 1 studies jointly and the power parameter of the last study.

Eq. 24 can be generalized, to reflect a situation where there are *P* disjoint sets of studies, denoted by 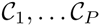, with genetic correlation within each set, but no genetic correlation between the sets. In this scenario, the power parameter *d* in Eq. 21 for a joint meta-analysis of all sets is given by

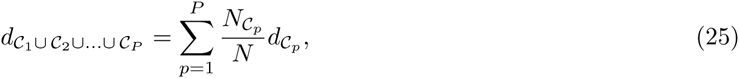

where 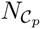 denotes the total sample size in study-set *C*_*p*_ and 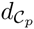 the power parameter in Eq. 21 for the meta-analysis of all studies in set *C*_*p*_, and *N* the total sample size when aggregating over all study sets. This equation states that power parameter for a meta-analysis of several sets of studies with CGR within each set, but no CGR between sets, is a weighted average of the power parameters in the underlying sets.

Since the statistical power is a monotonically increasing function of the power parameter *d*, Eq. 25 leads to two corollaries under CGR equal to zero between sets of studies, namely that

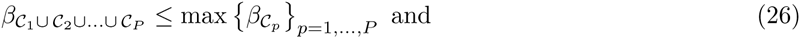

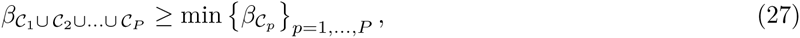

where 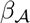 denotes the power in set of studies 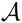.

The implication of Eq. 25 is simple yet powerful; when several sets of studies with genetic correlation within each set, but no genetic correlation between sets, are considered for meta-analysis, one should not meta-analyze sets 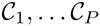 jointly, but rather meta-analyze only the set of studies which has the largest power parameter according to Eq. 21.

Only when 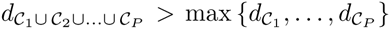, does the meta-analysis of all sets jointly have higher statistical power than a meta-analysis based on only one set of studies.

### S2 Derivations Accuracy

This section extends the theoretical framework for meta-analytic power. Derivations are based on the same assumptions as in S1 Derivations Power. We consider the predictive accuracy of the polygenic score (PGS) including all *S* independent SNPs, with SNP-weights based on the meta-analysis results from the set of *C* study, in a hold-out sample indexed as ‘study’ *C* + 1. In this hold-out sample, we focus exclusively on the theoretical *R*^2^ of the PGS; instead of considering *N*_*C*+1_ realizations of the stochastic processes underlying the genotypes and treating these as fixed explanatory variables, we treat the phenotype, the PGS, and the underlying genotypes as random variables, and use probability theory to derive *R*^2^. The hold-out sample is also allowed a study-specific SNP-based heritability, 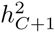, and genetic-correlations with the other *C* studies (thus extending both the CGR matrix and its Cholesky decomposition to (*C* + 1) × (*C* + 1) matrices).

First, we write the phenotype in hold-out sample as a function of noise and the independent genetic factors discussed in the preceding section. Second, we derive an expression for the PGS as a function of the genetic factors. Third, using this representation we derive the theoretical covariance between the PGS and the phenotype. Fourth, using the theoretical variances and covariance, we obtain an expression for the theoretical *R*^2^.

#### Polygenic model

Here, we derive an expression for the phenotype in the hold-out study as a function of independent genetic factors and an expression for the phenotypic variance.

Aggregating across causal SNP set 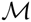 and the noise, the phenotype in study *C* + 1 can be written as follows:

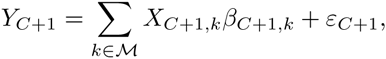

where, analogous to Eq. 12,

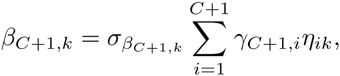

where *η*_*ik*_ now indicates the *i*-th element of the now (*C* + 1)-dimensional vector of independent normal draws, ***η***_***k***_, and where *γ*_*C* + 1,*i*_ describes an element of the Cholesky decomposition **Γ**_**G**_ of the (*C* + 1) × (*C* + 1) cross-study genetic correlation matrix, incorporating the hold-out sample. Hence, the phenotype can be written as

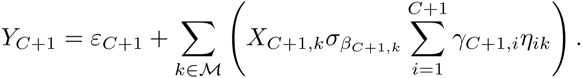

Analogous to the scaling of SNPs in S1 Derivations Power here, with genotypes treated as random variables, we assume

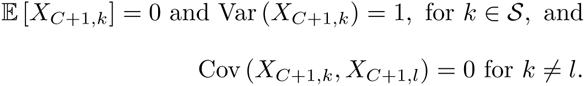

Consequently, the phenotypic variance in the hold-out sample is given by

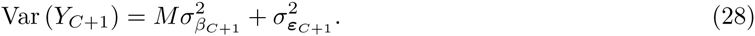

#### Polygenic score

Here, we derive an expression for the PGS as a function of independent genetic factors, an expression for the PGS variance, and its covariance with the phenotype in the hold-out sample.

Since each SNP in each study in the meta-analysis has been scaled such that its dot product equals the sample size of that study, by analogy of the standard error of the SNP effect estimate in a single study, the standard-error of the meta-analytic effect estimate 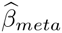 for study *C* + 1 can be approximated by

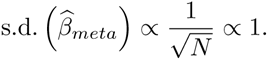

Hence, the meta-analytic effect estimate is proportional to the meta-analysis *Z* statistic. Since any scalar multiple of the PGS will not affect its *R*^2^ with respect to the phenotype, the *Z* statistics of the meta-analysis can be applied as SNP weights directly. Therefore, the PGS in the hold-out sample, including all SNPs, is given by

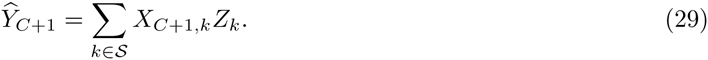

Plugging the expression for *Z*_*k*_ from Eq. 16 into Eq. 29, and substitution of terms by means of the square root of Eq. 20, the PGS is given by

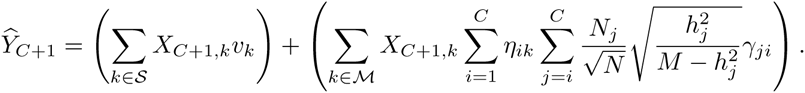

Exploiting the fact that *η*_*ik*_, υ_*k*_, and *X*_*C*+ 1,*k*_ are all independent random variables, with mean zero and variance one, we find that the variance of the PGS is given by

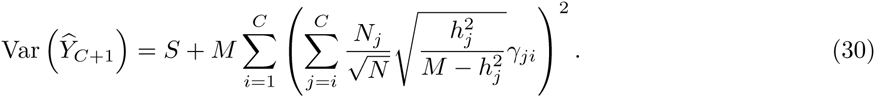

Again exploiting independence, zero mean, and unit variance of the respective terms, the covariance between the PGS and the phenotype is given by

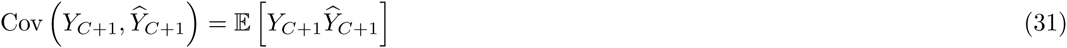

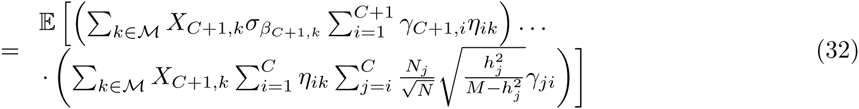

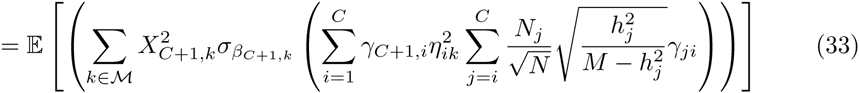

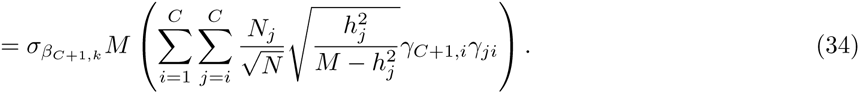

#### Theoretical *R*^2^

Here, we derive the theoretical *R*^2^ between the PGS and the phenotype in a hold-out study. For intuition, we present the theoretical *R*^2^ for a scenario with one study for discovery and one study as hold-out sample.

By combining Eq. 28, 30, and 34, the *R*^2^, defined as the squared correlation of the outcome and the PGS in the hold-out sample, is now given by

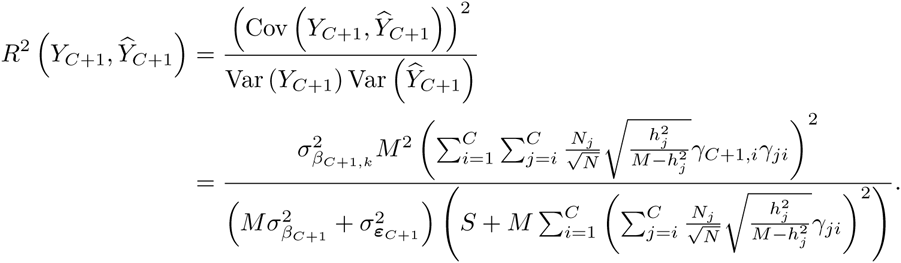

This expression can be simplified as follows:

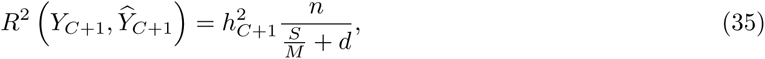

where *d* is the meta-analysis power parameter given in Eq. 21 and numerator *n* is given by

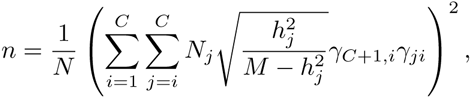

where *N* is the total sample size in the meta-analysis.

The expression for *R*^2^ in Eq. 35 is such that, in addition to the parameters needed for the power calculation, one only needs the genetic correlation between the hold-out sample and the meta-analysis samples and the heritability in the hold-out sample.

In case the number of studies for discovery is one (i.e., *C*=1), with a total sample size *N*, and with a genetic correlation *ρ*_G_ between the hold-out and discovery sample, we have that

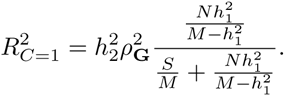

As in S1 Derivations Power, we have that under high polygenicity 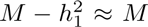. Therefore, an easy approximation of *R*^2^ in this scenario is given by

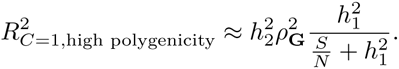

When 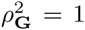, *S*=*M*, and 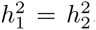, we obtain a known expression for PGS *R*^2^ in terms of sample size, heritability, and the number of SNPs [25]. In case 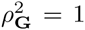 and we consider the *R*^2^ between the PGS and genetic value (i.e., the genetic component of the phenotype), both 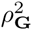 and 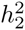 can be ignored, thereby making the last expression equivalent to the first equation in [37].

### S3 Note on Genetic Correlations

Consider, without loss of generality, a model for two phenotypes, *y*_1_ and *y*_2_. In line with Assumption 5 in S1 Derivations Power, let each causal variant, for the phenotype of interest, have the same *R*^2^ with respect to that phenotype.

In line with the random effects model, adopted in S1 Derivations Power and S2 Derivations Accuracy, we can write the data-generating processes of the respective phenotypes as

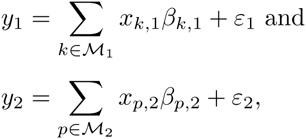

where 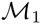 (resp. 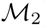) denotes the set of causal SNPs for *y*_1_ (*y*_2_) and where *β*_*k*,1_ (resp. *β*_*p*, 2_ the effect of *x*_*k*,1_ (*x*_*p*,2_), that is, standardized SNP *k* (p), on phenotype 1 (2).

The genetic correlation at the genome-wide level can now be conceptualized as the correlation in the true genetic value for both phenotypes. That is

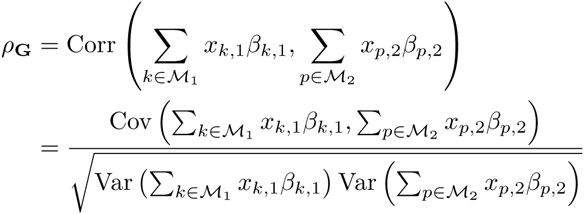

Assuming independent haplotype blocks with independent effects (Assumption 4), where the effects have mean zero, this expression for the genetic correlation at the genome-wide level can be rewritten as

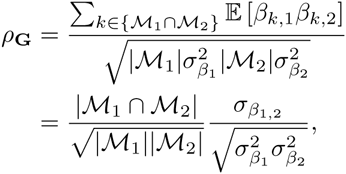

where 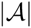 denotes the number of elements in set 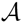.

Hence, the genetic correlation at the genome-wide level can be written as the product of overlap in causal loci between the two traits and the cross-trait correlation of the effects of these overlapping loci. That is,

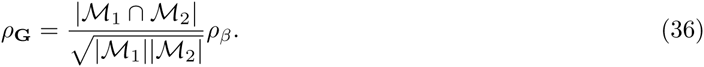

Eq. 36 is a generalization of the ‘common-elements formula’ [50], describing a correlation as a function of the number of overlapping elements and unique elements.

In particular, when 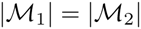, we have that

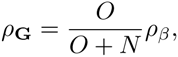

where *O* denotes the number of overlapping causal loci and *N* the number of idiosyncratic causal loci per trait.

We assume throughout the paper that all causal loci are shared across traits and studies (Assumption 3 in S1 Derivations Power). That is,

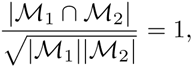

and that, consequently, the genetic correlation at the genome-wide level is equal to the correlation in the effects of overlapping causal SNPs. That is,

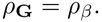

As we show in S4 Simulation Studies, the theoretical predictions of GWAS power and predictive accuracy obtained under this assumption are quite accurate, even when an imperfect genetic correlation at the genome-wide level is shaped primarily by lack of overlap in causal loci, rather than a poor correlation in the effects of overlapping loci.

### S4 Simulation Studies

Using five simulation studies, we assess the accuracy of the MetaGAP calculator, which is based on the expressions for GWAS power and PGS *R*^2^ derived in S1 Derivations Power and S2 Derivations Accuracy. Since the calculator is based on specific assumptions regarding the data-generating process, an important question is whether the calculator still provides accurate predictions of power and *R*^2^ when the underlying assumptions are violated.

Hence, each simulation study has a different underlying data-generating process. The first study, Simulation 1, assumes that rare variants have larger effects than common variants to such an extent that each causal SNP, regardless of allele frequency, is expected to have the same *R*^2^ with respect to the phenotype (Assumption 5 in S1 Derivations Power). This simulation is entirely in line with the assumptions underlying the MetaGAP calculator. In the second study, Simulation 2, common variants have effects of the same magnitude as rare variants (leading a common causal variant to explain a larger proportion of the phenotypic variation that a rare causal variant). The third study, Simulation 3, also allows for differential *R*^2^ between SNPs and, in addition, does not assume that SNP allele frequencies are uniformly distributed. Instead, the third study assumes that there are more variants in the lower minor allele frequency bins than in the higher minor allele frequency bins. In addition to the deviations from assumptions made in Simulations 2 and 3, Simulation 4 allows allele frequencies to be completely independent across studies. Finally, in Simulation 5, we go back to a data-generating process in line with the assumptions underlying the MetaGAP calculator, with one important difference; in Simulation 5, the genetic correlation as inferred at the genome-wide level is not only shaped by the correlation of SNP effects, but also by the degree of overlap of causal loci across studies. Thereby, Simulation 5 violates the assumption discussed in S3 Note on Genetic Correlations, that the estimated CGR is shaped only by imperfect correlations of SNP effects across studies.

For each simulation study there are 100 independent runs. In each run data is simulated for *C*=3 distinct samples for discovery as well as a fourth sample used as hold-out sample for prediction. The sample sizes of the respective studies are given by *N*_1_=20, 000, *N*_2_=15, 000, *N*_3_=10, 000, and *N*_4_=1, 000, where *N*_4_ denotes the sample size of the hold-out sample. For Simulations 1-4, an 11 × 11 grid of equispaced values of 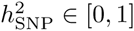 and *ρβ* ∈ [0, 1] is considered. Similarly, for Simulation 5, an 11 × 11 grid of equispaced values of *s* ∈ [0, 1] and *ρβ* ∈ [0, 1] is considered. Here, *s* denotes the fraction of causal SNPs that overlaps across studies and *ρβ* the cross-study correlation of the effects of SNPs that are overlapping. In Simulations 1–4 we have that *s*=1 and in Simulation 5 we have that 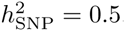. In all simulations there are *S*=100, 000 independent SNPs of which *M*=1, 000 have a causal influence. Moreover, when computing theoretical power and predictive accuracy, in line with S3 Note on Genetic Correlations, we use *ρ*_G_=*s* · *ρ*_G_ as value of the input parameter CGR. A detailed description of the data-generating process in each simulation study can be found in Table 3.

**Table 3.**
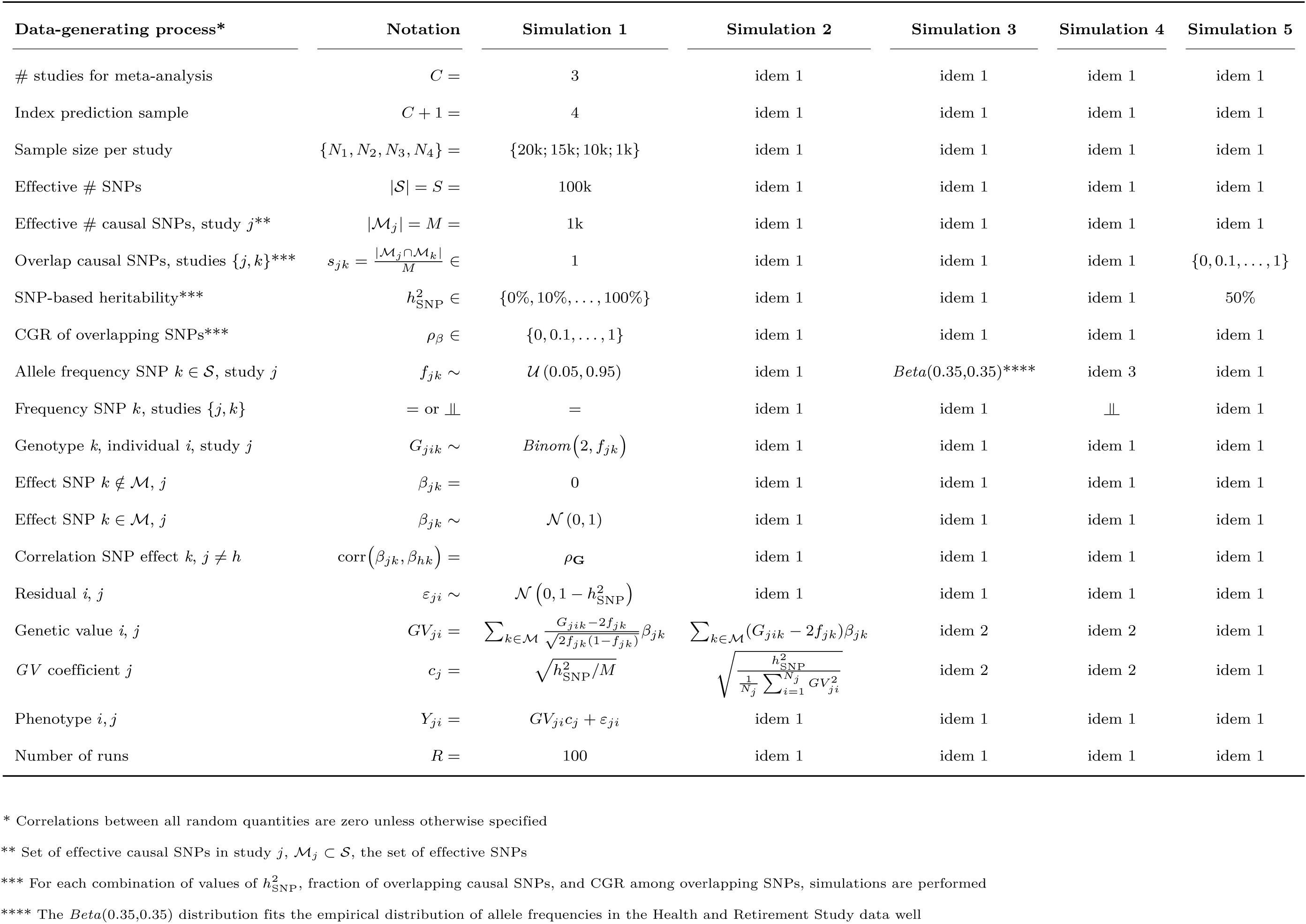
Design of Simulations 1–5. Design settings identical to a preceding simulation study are denoted by ‘idem’, followed by the number of the first preceding simulation study with the same setting.

For every run, data is simulated in accordance with the underlying data-generating process. Next, a GWAS is carried out in each of the three discovery samples. GWAS results are then meta-analyzed using sample-size weighting. The fraction of causal SNPs reaching genome-wide significance in the meta-analysis is the estimate of statistical power per SNP. The squared correlation between the meta-analysis-based PGS for the hold-out sample and the corresponding phenotype is the estimate of the PGS *R*^2^.

Final estimates of power per causal SNPs and PGS *R*^2^ are obtained by averaging the estimates across the runs. Fig. 6–7, show the resulting estimates of power per causal SNP in the meta-analysis and the *R*^2^ of the PGS, for both Simulations 1–4 and Simulation 5. In addition, both figures report the power per causal SNP and *R*^2^ predicted by the theoretical model, derived under the assumptions discussed in S1 Derivations Power. Inspection of Fig. 6 shows that there is no qualitative difference between the contour plots. Moreover, when computing the root-mean-square error (RMSE) between the theoretical predictions and the simulation-based estimates of power and *R*^2^, even for the most extreme departures from our assumptions regarding allele frequencies and effects sizes (Simulations 3–4), the RMSE in power remains below 3% and the RMSE in *R*^2^ of the PGS below 2%. Hence, the theoretical predictions of GWAS power and predictive accuracy – derived under assumptions of equal true *R*^2^ of causal SNPs, with uniformly distributed allele frequencies that are equal across studies – are robust to violations of these assumptions.

**Figure 6.**
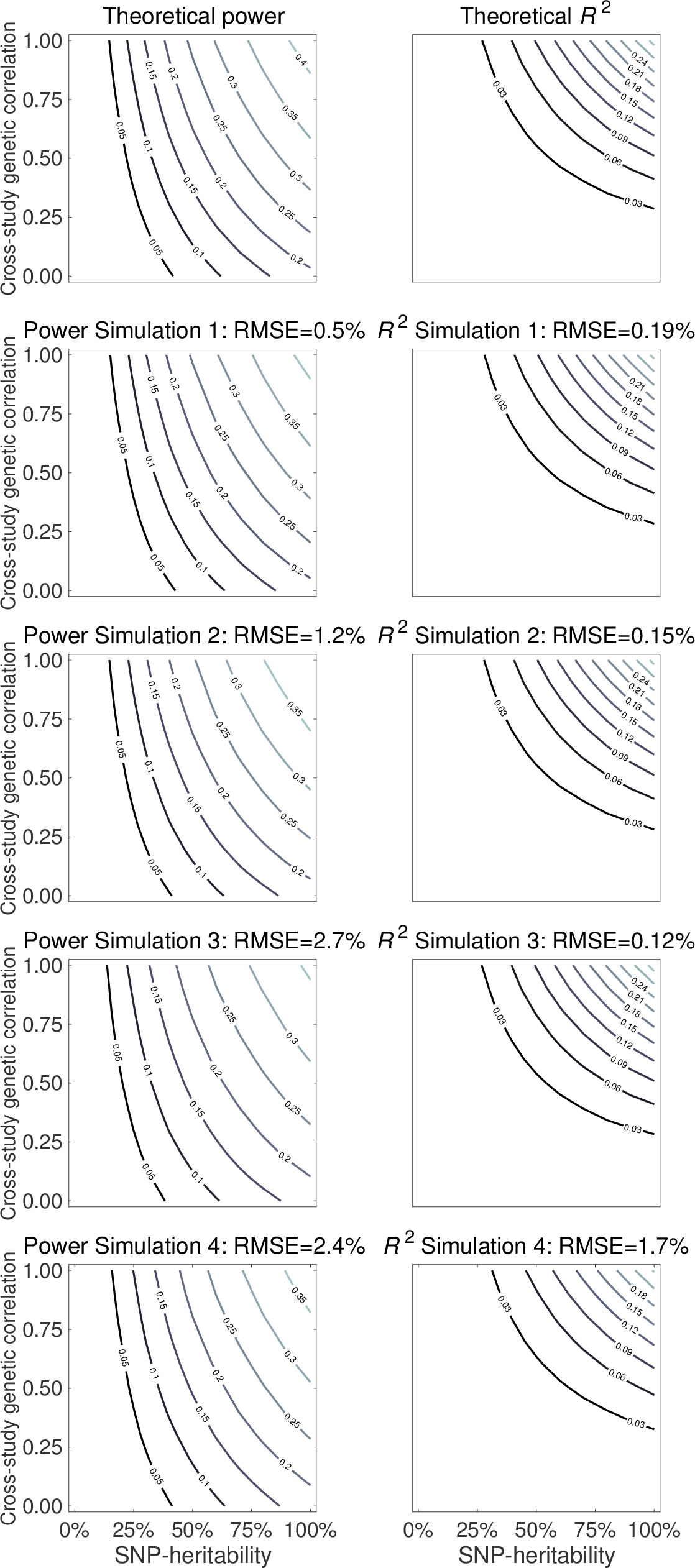
Power and polygenic score *R*^2^ contour plots, with in each plot h^2^ on the *x*-axis and cross-study genetic correlation on the *y*-axis. The first row shows predictions from the theoretical model. Subsequent rows show estimates based on respective simulation studies. The first column shows power per causal SNP. The second column the *R*^2^ of a polygenic score in a hold-out sample. Above each plot, the root-mean-square error (RMSE) is reported for the difference between predictions from the theoretical model and the simulation-based estimates.

**Figure 7.**
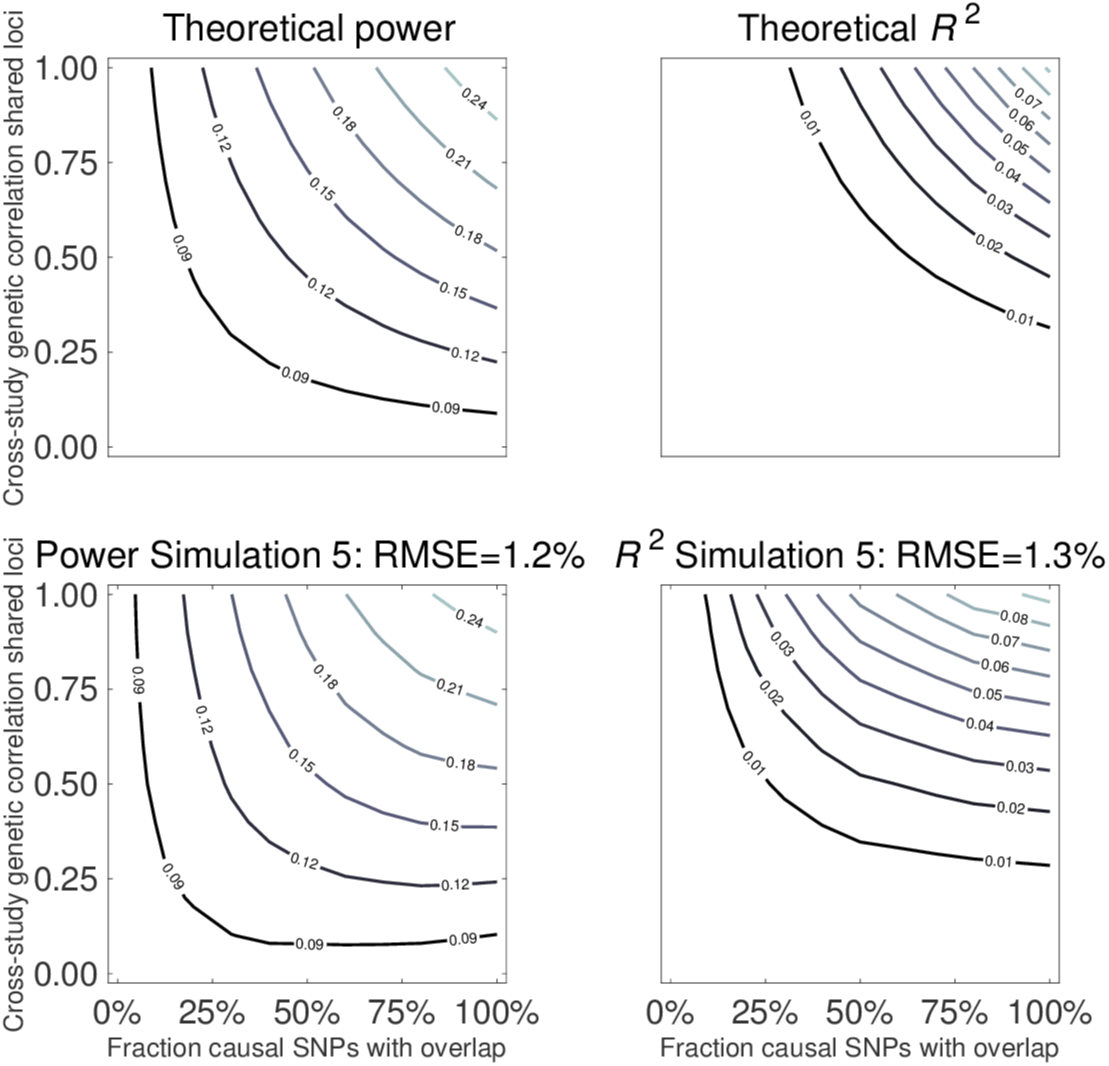
Power and polygenic score *R*^2^ contour plots, with in each plot the fraction of causal loci that overlaps across studies on the *x*-axis and cross-study correlation of the effects of overlapping loci on the *y*-axis. The first row shows predictions from the theoretical model. The second row shows estimates based on a simulation study. The first column shows power per causal SNP. The second column the *R*^2^ of a polygenic score in a hold-out sample. Above each plot, the root-mean-square error (RMSE) is reported for the difference between predictions from the theoretical model and the simulation-based estimates.

Inspection of Fig. 7 shows that when CGRs are being shaped by a combination of poor overlap and poorly correlated effects of overlapping loci, there are some qualitative differences between predicted power and predictive accuracy compared to simulation-based estimates. However, the RMSE of theoretical power is only 1.2% with respect to the power estimated from simulations. Similarly, the RMSE of theoretical predictive accuracy is only 1.3%. Hence, the quantitative differences are small.

Simulation 5 shows that when low CGRs are induced by poor overlap of causal loci across studies instead of low correlations of the effects of overlapping loci, this leads to a slight downwards bias in our theoretical predictions (i.e., making our theory conservative). Hence, we argue that if our calculator deems a study design well-powered, the analyses will be well powered, potentially even more so than what our theory predicts (e.g., if some of the imperfect CGR can be attributed to causal loci that are not shared across studies).

### S5 Data and Quality Control

#### Genotype data

In the bivariate and univariate genomic-relatedness-matrix restricted maximum likelihood (GREML) analyses we use genotype data from the Rotterdam Study (RS; Ergo waves 1-4 sample denoted by RS-I, Ergo Plus sample denoted by RS-II, and Ergo Jong sample denoted by RS-III), the Swedish Twin Registry (STR; TwinGene sample), and the Health and Retirement Study (HRS). For each study, details on the genotyping platform, quality control (QC) prior to imputation, the reference sample used for imputation, and imputation software, are listed in Table 4.

**Table 4.**
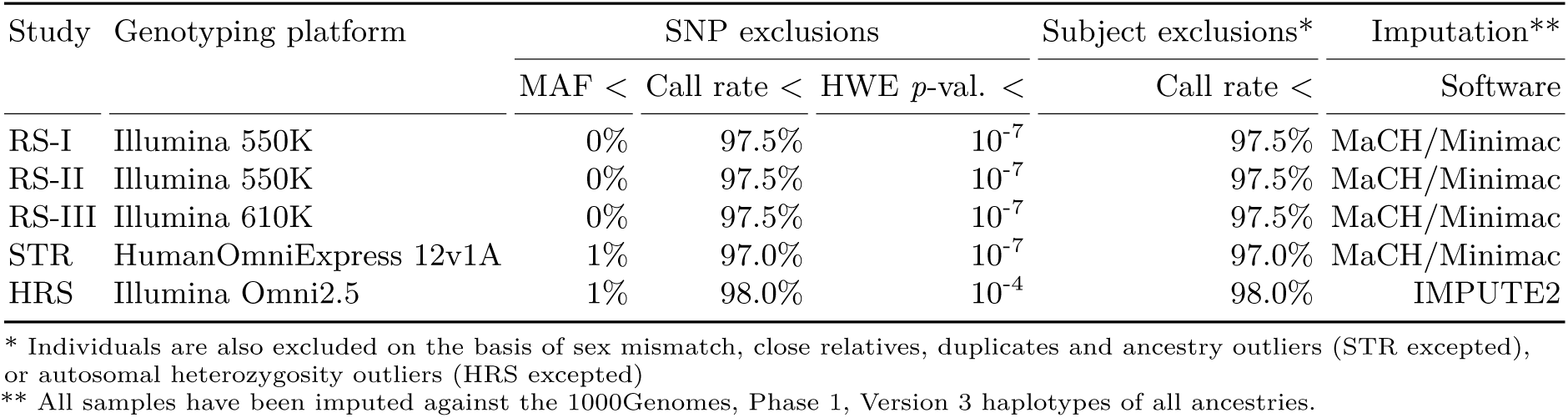
Genotyping and imputation.

To increase the overlap of SNPs across studies, we use genotypes imputed on the basis of the 1000 Genomes, Phase 1, Version 3 reference panel [51]. We only consider the subset of HapMap3 SNPs available in the 1kG data. By using this subset we substantially reduce the computational burden of the analyses, while preserving overlap between the SNP-sets in the studies and still having a sufficiently dense set of both common and more rare SNPs (# SNPs after QC ≈ 1 million).

#### Quality control

Prior to QC, we extract HapMap3 SNPs (source: http://hapmap.ncbi.nlm.nih.gov/downloads/genotypes/hapmap3_r3/plink_format/, accessed: December 11, 2014) from the imputed genotype data of each study and convert the allele dosages to best-guess `PLINK` [52, 53] binary files by rounding dosages using `GCTA` [35]. Subsequently, we perform QC on the best-guess genotypes in two stages. In the first stage, we clean and harmonize the imputed genotype data at the study level. The cleaned and harmonized study genotypes are then merged into a pooled dataset. The second round of QC is aimed at cleaning the pooled dataset, on the basis of the samples for which the phenotype is available. Hence, the first QC stage is phenotype-independent, whereas the second stage depends on the phenotype of interest.

In the first QC stage (prior to merging), we filter out the following markers and individuals:

1. SNPs with imputation accuracy below 70%.
2. Non-autosomal SNPs.
3. SNPs with minor-allele frequency below 1%.
4. SNPs with Hardy-Weinberg-Equilibrium *p*-value below 1%.
5. SNPs with missingness greater than 5%.
6. Individuals with missingness greater than 5%.
7. SNPs that are not present in all studies.
8. SNPs whose alleles cannot be aligned across studies.

Prior to the first QC stage, we apply the following two additional steps in HRS:

1. Switch alleles to address a strand-flip error due to incorrect annotation.
2. Drop individuals of non-European ancestry.

After the first round of QC, a set of roughly 1 million overlapping SNPs, available for about 30,000 individuals is left. Panel I in Table 5 shows, for each study, the number of SNPs and individuals before and after the first round of QC.

**Table 5.**
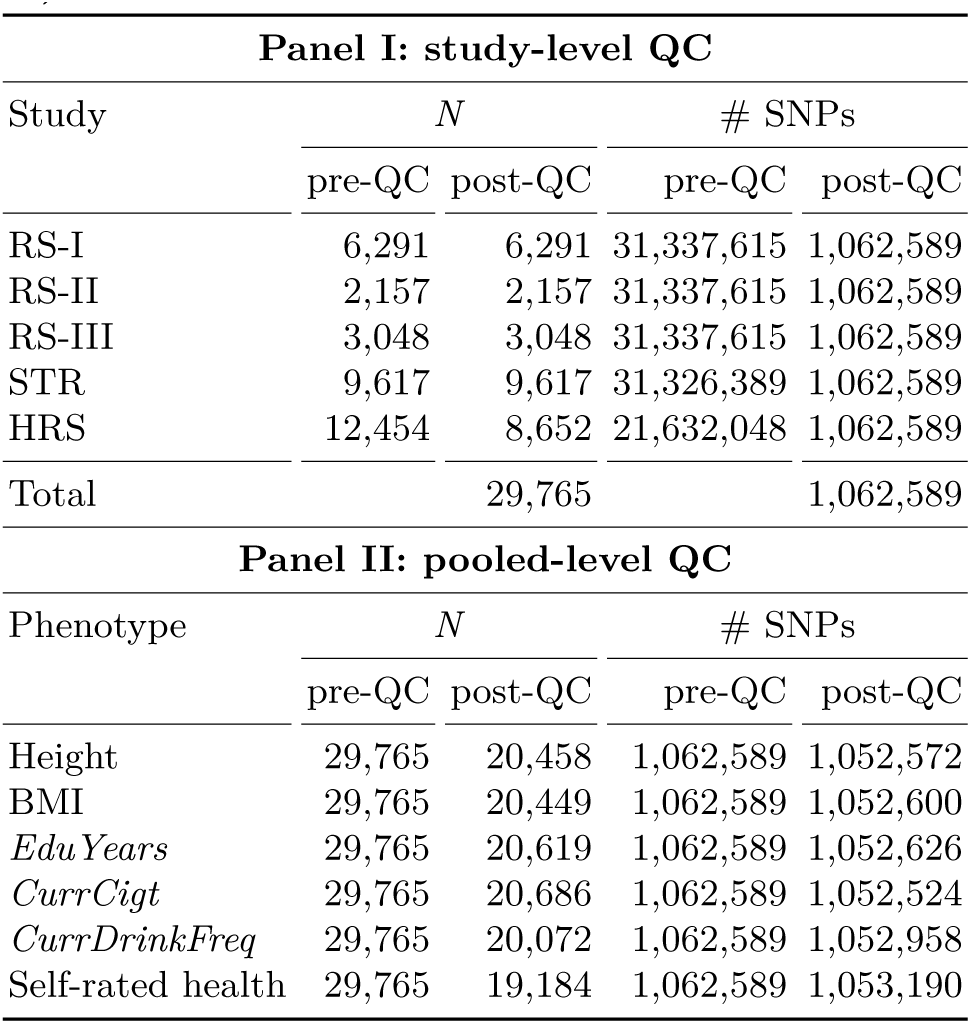
Number of individuals and SNPs before and after quality control (QC) at the study level (Panel I) and at the pooled level (Panel II).

The second QC stage, applied to the pooled data set, comprises the following steps:

1. Keep only individuals for whom the phenotype of interest and all corresponding control variables are available.
2. Drop SNPs with a minor-allele frequency below 1%.
3. Drop SNPs with Hardy-Weinberg-Equilibrium *p*-value below 1%.
4. Drop SNPs with missingness greater than 5%.
5. Drop individuals with missingness greater than 5%.
6. Keep only one individual per pair of individuals with a genomic relatedness greater than 0.025.

Since the data in STR consists of twins and having highly related individuals can bias estimates of SNP-based heritability due to environment-sharing, we randomly select only one individual per twin pair after Step 1 in the second QC stage.

Panel II in Table 5 shows the sample size and the number of SNPs in the pooled dataset for each phenotype. We only consider phenotypes that attain a sample size of at least 18,000 individuals after all QC steps. The lowest sample size after QC is 19,184 for self-rated-health and the highest is 20,686 for *CurrCigt*. For all phenotypes, the number of SNPs is slightly greater than one million.

#### Phenotype data

For HRS, we use the RAND HRS data, version N, to obtain the phenotypes of interest. These data consist of measurements from eleven waves. RS-I consists of four data waves (Ergo 1-4). In both HRS and RS-I, data for some phenotypes are only available in a subset of the waves. RS-II, RS-III and STR do not have multiple measures over time for the phenotypes considered in this study. Table 6 describes how the phenotypes are constructed in each of the five studies.

**Table 6.**
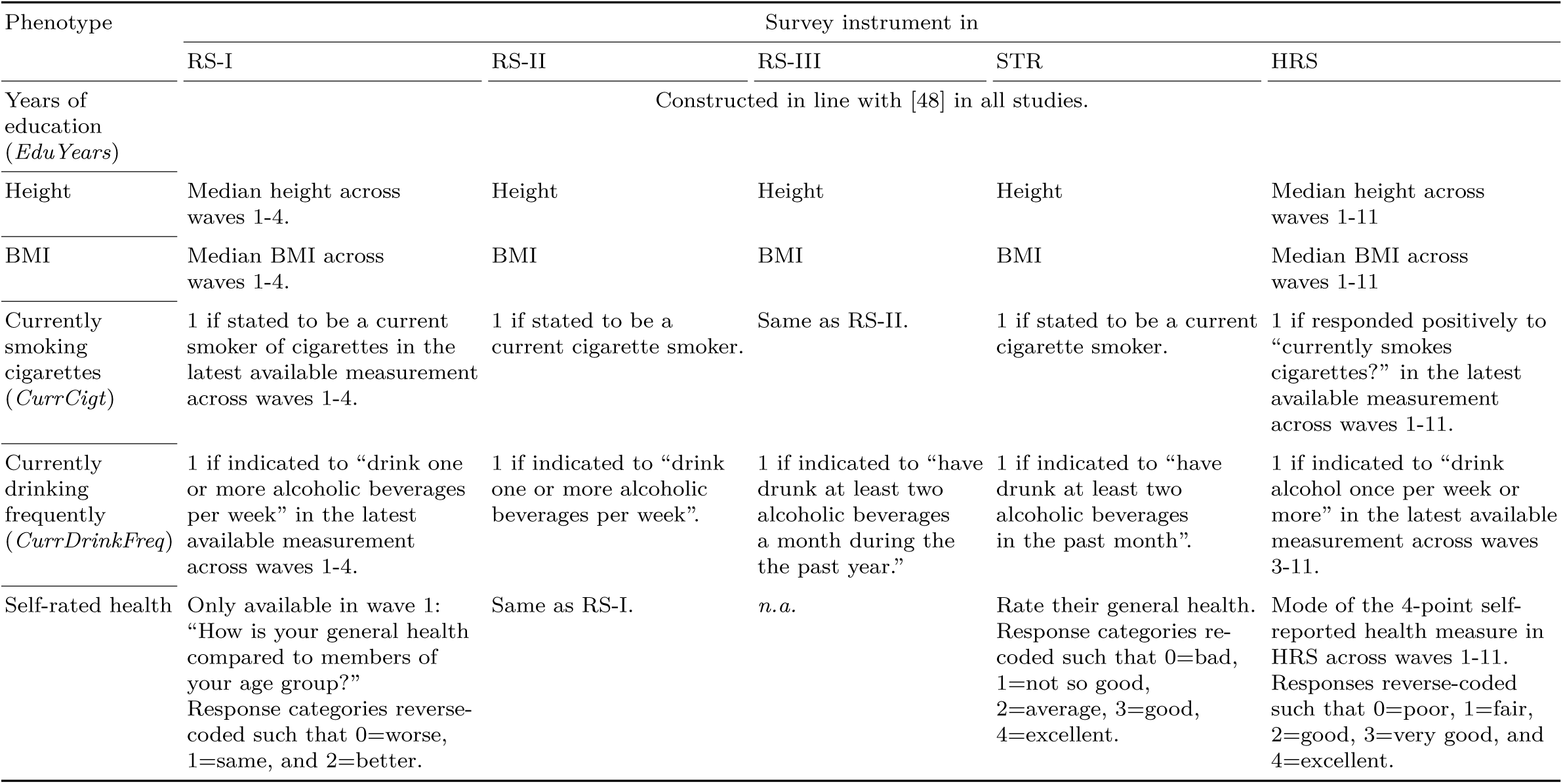
Study-level phenotype measures.

As Table 6 shows, height, BMI, *EduYears*, and *CurrCigt* are measured quite consistently across waves. The self-rated health phenotype is also measured quite consistently, although in RS respondents are asked about health compared to members of the same age group, whereas a more absolute question is posed in STR and HRS. The drinking measure *CurrFreqDrink* is also measured somewhat heterogeneously; the threshold for what we treat as ‘frequent drinking’ is determined by how fine-grained the drinking frequency measure is in the respective studies.

### S6 GREML Estimation

Height, BMI, *EduYears*, and self-rated health are treated as quantitative traits. *CurrCigt* and *CurrDrinkFreq* are treated as binary outcomes. In each study, (after aggregating across waves, if applicable) we regress quantitative phenotypes on age, squared age, sex, and an intercept. The residuals from the regression are standardized to have a sample-mean equal to zero and variance equal to one. For both binary and quantitative traits, the aforementioned covariates are also included in the GREML estimation. In addition, in bivariate GREML and pooled GREML estimation (i.e., considering multiple studies jointly), the intercept is replaced by indicator variables for the respective studies, capturing study-specific fixed effects. Finally, 20 principal components from the phenotype-specific genomic-relatedness matrix are added to the set of control variables in the GREML estimation, in order to correct for population stratification [54].

### S7 GREML Results

Details per phenotype on sample size, univariate estimates of SNP heritability, and bivariate estimates of genetic correlation, stratified across studies, and cross-study averages, are provided in Table 7. Results stratified across sexes are listed in Table 8.

**Table 7.**
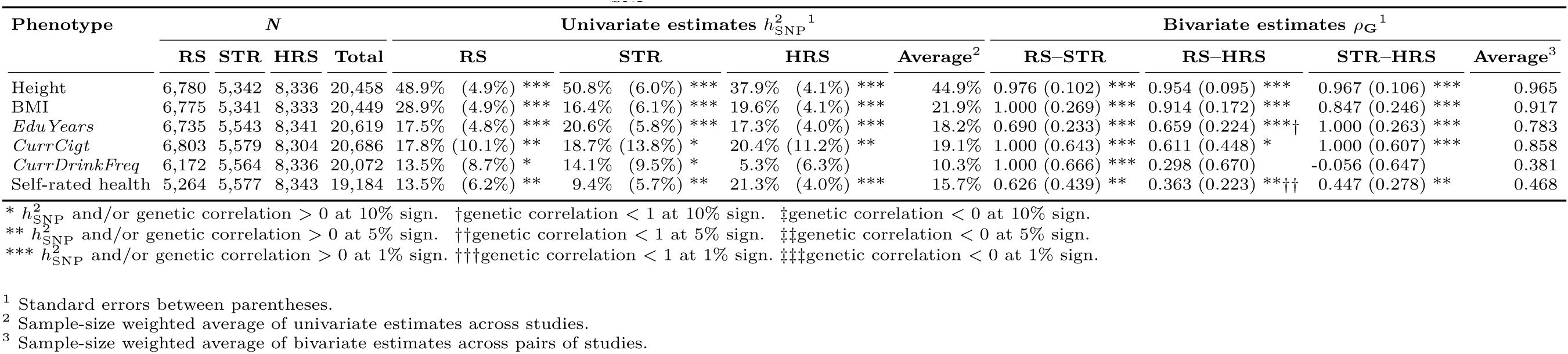
GREML estimates of SNP heritability (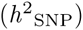) and genetic correlation (*ρ*_G_) across studies.

**Table 8.**
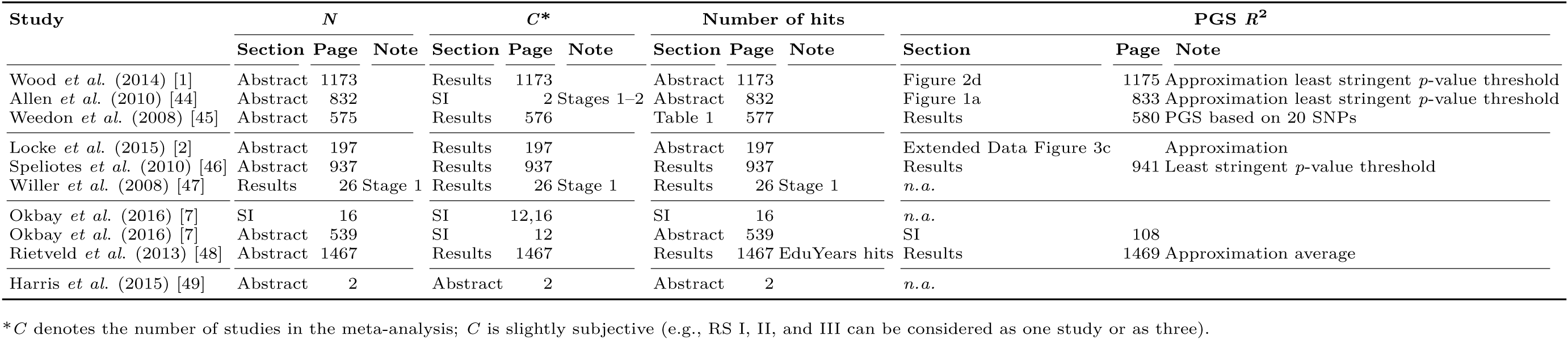
GREML estimates of SNP heritability (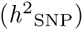) and genetic correlation (*ρ*_G_) across sexes.

### S8 Large-scale GWAS efforts

Table 9 shows the meta-analysis packages, and the assumptions underlying those packages, used in large-scale GWAS efforts for the traits considered in our attenuation study, reported in Table 2. Similarly, Table 10 shows details and notes on the results from large-scale GWAS efforts that are used as input in the aforementioned attenuation study.

**Table 9.**
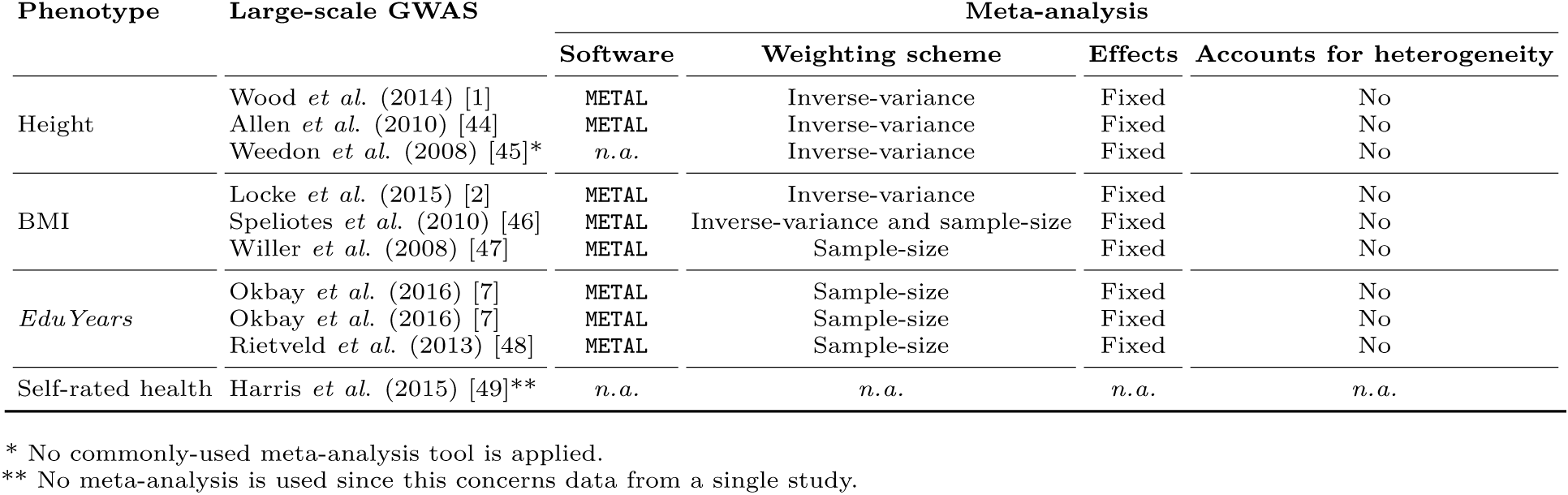
Meta-analysis methods used in large-scale GWAS efforts to date for height, BMI, *EduYears*, and self-rated health, reported in the same order as in Table 2.

**Table 10.**
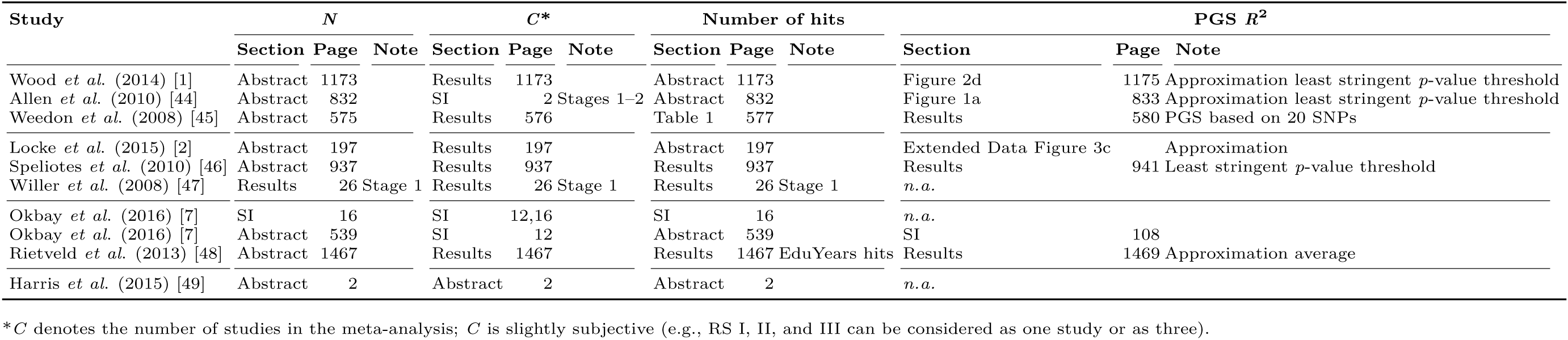
Details and notes on the sources for the sample size, the number of meta-analyzed studies, the number of genome-wide-significant hits, and the PGS *R*^2^, for large-scale GWAS efforts to date for height, BMI, *EduYears*, and self-rated health, reported in the same order as in Table 2.

## Acknowledgments

**RS (Rotterdam Study)** The generation and management of GWAS genotype data for the Rotterdam Study is supported by the Netherlands Organisation of Scientific Research NWO Investments (nr. 175.010.2005.011, 911-03-012). This study is funded by the Research Institute for Diseases in the Elderly (014-93-015; RIDE2), the Netherlands Genomics Initiative (NGI)/Netherlands Organisation for Scientific Research (NWO) project nr. 050-060-810. We thank Pascal Arp, Mila Jhamai, Marijn Verkerk, Lizbeth Herrera and Marjolein Peters for their help in creating the GWAS database, and Karol Estrada and Maksim V. Struchalin for their support in creation and analysis of imputed data. The Rotterdam Study is funded by Erasmus Medical Center and Erasmus University, Rotterdam, Netherlands Organization for the Health Research and Development (ZonMw), the Research Institute for Diseases in the Elderly (RIDE), the Ministry of Education, Culture and Science, the Ministry for Health, Welfare and Sports, the European Commission (DG XII), and the Municipality of Rotterdam. The authors are grateful to the study participants, the staff from the Rotterdam Study and the participating general practitioners and pharmacists. Researchers who wish to use data of the Rotterdam Study must obtain approval from the Rotterdam Study Management Team. They are advised to contact the PI of the Rotterdam Study, Dr. Albert Hofman.

**STR (Swedish Twin Registry)** The Jan Wallander and Tom Hedelius Foundation (P2012-0002:1), the Ragnar Söderberg Foundation (E9/11), The Swedish Research Council (421-2013-1061), the Ministry for Higher Education, The Swedish Research Council (M-2205-1112), GenomEUtwin (EU/QLRT-2001-01254; QLG2-CT-2002-01254), NIH DK U01-066134, The Swedish Foundation for Strategic Research (SSF). Researchers interested in using STR data must obtain approval from the Swedish Ethical Review Board and from the Steering Committee of the Swedish Twin Registry. Researchers using the data are required to follow the terms of an Assistance Agreement containing a number of clauses designed to ensure protection of privacy and compliance with relevant laws. For further information, contact Patrik Magnusson.

**HRS (Health and Retirement Study)** The HRS (Health and Retirement Study) is sponsored by the National Institute on Aging (grant number NIA U01AG009740) and is conducted by the University of Michigan. The genotyping was funded separately by the National Institute on Aging (RC2 AG036495, RC4 AG039029). Our genotyping was conducted by the NIH Center for Inherited Disease Research (CIDR) at Johns Hopkins University. Genotyping quality control and final preparation of the data were performed by the Genetics Coordinating Center at the University of Washington. Genotype data can be accessed via the database of Genotypes and Phenotypes (dbGaP, http://www.ncbi.nlm.nih.gov/gap, accession number phs000428.v1.p1). Researchers who wish to link genetic data with other HRS measures that are not in dbGaP, such as educational attainment, must apply for access from HRS. See the HRS website (http://hrsonline.isr.umich.edu/gwas) for details.

**RAND HRS** RAND HRS Data, Version N. Produced by the RAND Center for the Study of Aging, with funding from the National Institute on Aging and the Social Security Administration. Santa Monica, CA (September 2014). Researchers who wish to use the RAND HRS data need to register via the RAND website (http://www.rand.org/labor/aging/dataprod/hrs-data.html).

**Dutch national e-infrastructure** This work was carried out on the Dutch national e-infrastructure with the support of SURF Cooperative.

